# TrackSOM: mapping immune response dynamics through sequential clustering of time- and disease-course single-cell cytometry data

**DOI:** 10.1101/2021.06.08.447468

**Authors:** Givanna H. Putri, Jonathan Chung, Davis N. Edwards, Felix Marsh-Wakefield, Suat Dervish, Irena Koprinska, Nicholas J.C. King, Thomas M. Ashhurst, Mark N. Read

## Abstract

Mapping the dynamics of immune cell populations over time or disease-course is key to understanding immunopathogenesis and devising putative interventions. We present TrackSOM, an algorithm which delineates cellular populations and tracks their development over a time- or disease-course of cytometry datasets. We demonstrate TrackSOM-enabled elucidation of the immune response to West Nile Virus infection in mice, uncovering heterogeneous sub-populations of immune cells and relating their functional evolution to disease severity. TrackSOM is easy to use, encompasses few parameters, is quick to execute, and enables an integrative and dynamic overview of the immune system kinetics that underlie disease progression and/or resolution.

## 1 Background

The immune response is a dynamic process in time, and inter-individual differences therein often underlie diverging clinical outcomes. Under challenge, the immune response deviates from homeostasis with the expansion of numerous cellular phenotypes that enact challenge-specific effector functions in a temporally coordinated fashion. This is seen, for instance, with accelerated cellular differentiation in the bone marrow and the infiltration of immune cell populations into peripheral tissues. Yet, the immune response is fallible, as attested to by autoimmunity, chronic diseases and lethal infections, all of which can be mediated by the immune response and may be ameliorated by immune intervention. Greater understanding of the temporal kinetics and interactions of immune cell populations would present expanded opportunities to intervene for clinical benefit. Essential to delivering this vision is an explicit mapping of how immune system status varies with time (‘*temporal mapping*’) and how that in turn associates with disease severity.

Cytometry is a popular and widely accessible technology for phenotypically characterising and quantifying immune cells from a sample (e.g. blood), and taking samples over time or disease progression presents an opportunity for developing such mappings of immune response against disease stage. Cytometry quantifies numerous characteristics (‘*markers*’) on potentially millions of individual cells. Advancements in this technology have now enabled measurements of upwards of 45 markers per cell, and this will continue to grow (Bendall *et al*., 2014; Park *et al*., 2020; Mair and Prlic, 2018; Nettey *et al*., 2018). A key analytical challenge in cytometry is aggregating (‘*gating*’) these many and heterogeneous cells into their distinct cellular phenotypes. Given the high-dimensional nature of such data, the task is challenging, requiring considerable domain expertise, whilst being prone to subjectivity and poor reproducibility (Mair *et al*., 2016; Maecker *et al*., 2012; Saeys *et al*., 2016). Many automated ‘clustering’ algorithms have emerged to support this activity (Liu *et al*., 2020; Qiu *et al*., 2011; Van Gassen *et al*., 2015; Levine *et al*., 2015). Of these, FlowSOM has gained popularity, being simple to use, fast to execute, and outstanding in reproducing manual-gating efforts (Putri *et al*., 2021; Weber and Robinson, 2016).

Lacking, however, is algorithmic support for annotating the immune response dynamics with respect to time and disease stage. This requires both (i) clustering to identify populations in each dataset in the time/disease-stage sequence, and (ii) tracking how these populations evolve, which can be complex. Both the absolute number of cells for each phenotype, and their relative representation within the sample, are liable to change. So too are population marker expression levels, which change as populations undergo differentiation, maturation and associated functional alteration. Furthermore, differentiation, such as from haematopoietic stem cells through common progenitors into distinct effector phenotypes, mean that phenotype trajectories through marker-space can present as branching tree-like structures through the time/disease-stage sequence. Infiltration into and egress out of peripheral tissues can manifest as populations appearing and disappearing. Identifying the existence and timing of phenotypic sub-populations that can impact on disease status is also highly relevant (Krzywinska *et al*., 2016). Required is a unified clustering-and-tracking algorithmic framework that can deliver these insights. Filling this knowledge gap, we previously developed ChronoClust (Putri *et al*., 2019a), a clustering algorithm wherein the data themselves determine the number and shape (in marker space) of immune cell populations. Importantly, ChronoClust can track the evolution of clusters across a sequence of datasets using a tracking mechanism which (i) requires no tunable hyperparameters, and (ii) is independent of the clustering algorithm used and can thus be ported onto other algorithms. Whilst ChronoClust’s clustering performance is highly tunable, it is computationally cumbersome and finding appropriate parameters can prove challenging, whereas we have found FlowSOM to offer better clustering performance (Putri *et al*., 2021).

Here, we present TrackSOM, a novel automated clustering and phenotype-tracking algorithm for cytometry data. It combines the high-quality clustering capacity and fast run time of FlowSOM with the tracking capability of ChronoClust enabling it to map immune cell population dynamics against time and/or disease severity status. We highlight these capabilities on a synthetic dataset and by recapitulating previous findings on a West Nile Virus (WNV) bone marrow dataset. We perform a parameter sensitivity analysis and demonstrate TrackSOM to have both an improved clustering performance and lower sensitivity to parameter value selections over ChronoClust. We offer usage and parameter selection advice to potential users with novel data. Lastly, we demonstrate usage on novel data by characterising the evolving immune response in the brains of WNV-infected mice and relating this to disease severity. TrackSOM is open-source and available at https://github.com/ghar1821/TrackSOM.

## 2 Results

### 2.1 The TrackSOM algorithm

TrackSOM takes as input a sequence of datasets, where each dataset captures cytometric quantifications on a collection of cells, and the sequence represents time-points or disease-stages, Figure 1A. TrackSOM determines clusters, representing phenotypes, within each dataset and then establishes links between them to reveal temporal evolutions between clusters across the sequence. TrackSOM extends FlowSOM in providing this functionality. A self-organising map (SOM) is built upon the amalgamated data from all datasets, thus identifying the data’s broad topological shape across the entire sequence, Figure 1B. Each data-point is uniquely associated with a single SOM node and the data that any given SOM node represents may be confined to a single or several time-point(s) or disease-stage(s). TrackSOM’s ability to track how clusters evolve is based on variation in which SOM nodes are supported by data-points as the dataset sequence progresses (Figure 1C) - some nodes will become more relevant, others less so.

**Figure 1:**
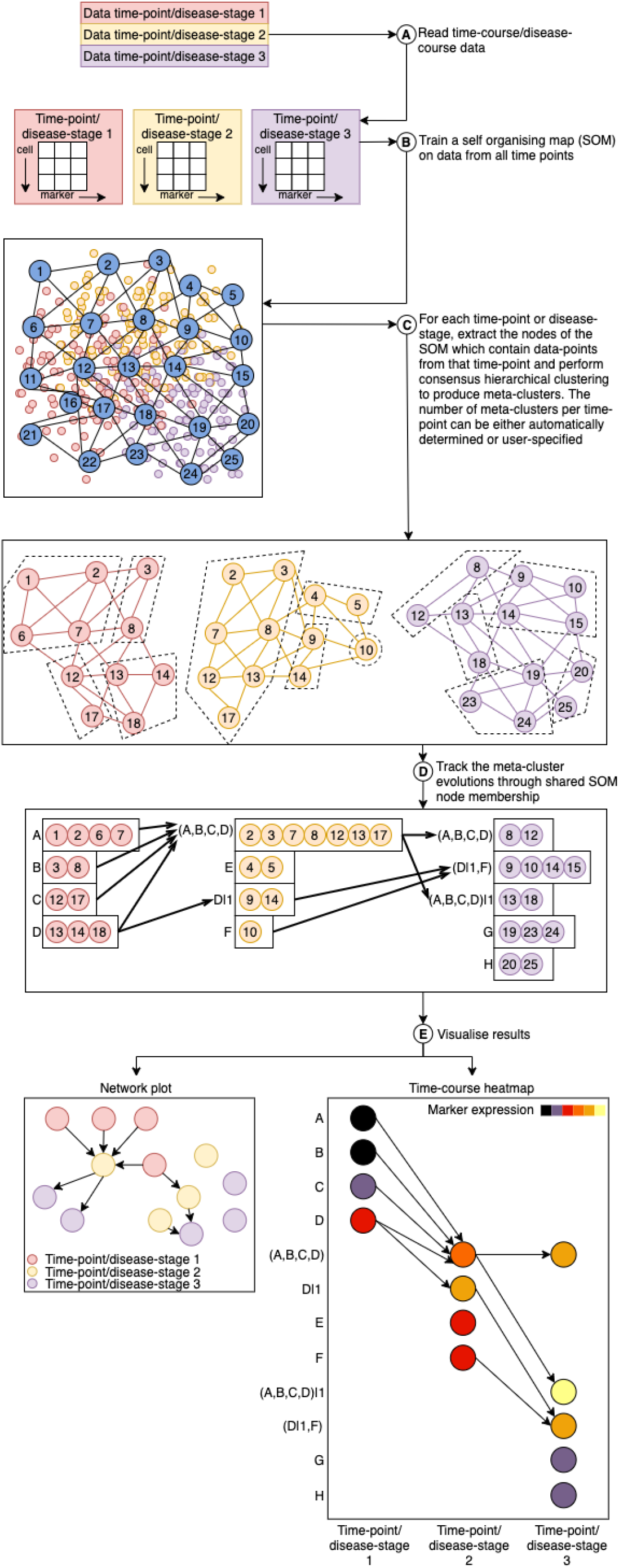
Schematic overview of TrackSOM’s operation and usage. (A) Time-course or disease-course cytometry data is parsed by TrackSOM, resulting in a matrix of cells (rows) and marker values (columns) per time-point/disease-stage. (B) A self-organising map (SOM) is then trained on all the matrices. Numbered circles represent SOM nodes. (C) SOM nodes containing data from each given time-point/disease-stage are then isolated and ‘meta-clustered’ (dotted polygons) using consensus hierarchical clustering. (D) The temporal evolutions of the resulting meta-clusters are tracked through their shared SOM node memberships. (E) Meta-cluster evolutions can be visualised as network plots (left) or time-course heatmaps (right). For clarity, the figure refers predominantly to ‘time-points’, but these could instead be datasets in a series of escalating disease severity. We have shown only 3 time-points, but TrackSOM is not limited to this.

Clusterings (phenotypes) of data-points at each time-point are captured through *meta-clusters*, which represent the collation of SOM nodes into discrete groupings, Figure 1C. Having built the SOM, TrackSOM then processes each dataset (time-point/disease-stage) independently, isolating SOM nodes containing data from that dataset only and forming meta-clusters accordingly. The centroids of these non-empty SOM nodes are re-calculated (temporarily disregarding data from other datasets) as per FlowSOM operation. Meta-clusters for the given dataset are derived through consensus as drawn from multiple efforts at sub-sampling and hierarchically clustering the SOM’s nodes, again in accordance with FlowSOM. The number of meta-clusters formed is parameter-defined, as discussed below.

Next, meta-cluster linkages across adjacent time-points/disease-stages are determined to track meta-cluster evolutions. This is accomplished by identifying those meta-clusters in adjacent datasets that share common SOM nodes, Figure 1D. This method is adapted from the ChronoClust algorithm (Putri *et al*., 2019a), and we direct readers there for full details. Briefly, meta-cluster evolutions are encoded into the meta-cluster IDs. *New meta-clusters* are defined as containing exclusively SOM nodes that did not belong to any meta-clusters in the preceding datasets. This includes the first time-point/disease-stage. New meta-clusters are assigned unused alphabetic labels from an alphabetical sequence (A, B,…, Z, AA, BB,…). Meta-cluster evolutions are captured in how these labels are amended over time. A meta-cluster that contains only a subset of a preceding meta-cluster’s SOM nodes is termed a *split meta-cluster*. A meta-cluster can simultaneously split multiple ways, and the child inheriting the majority of SOM nodes retains the parent’s label. The remaining sibling meta-clusters are assigned A|1, A|2, etc. A *merged meta-cluster* has captured SOM nodes belonging to multiple preceding meta-clusters, and its label is a parenthesised concatenation of its parent IDs, e.g. (A,B,C,D).

TrackSOM provides several visualisations, Figure 1E. *Network plots* denote all meta-clusters at all time points/disease-stages as bespoke nodes, linked via their evolutions. *Time-course heatmaps*capture time/disease-stage and meta-cluster IDs on separate axes. The visualisations are complementary: time-course heatmaps can become convoluted for temporally complicated cluster evolutions, and the network plot can provide a simpler global overview of the data. In both visualisations, node sizes indicate the proportion of a dataset’s data-points that the corresponding meta-cluster captures. Meta-clusters can be coloured by average marker expression-level to depict movements through marker-space as the sequence progresses, by time-point/disease-stage, or originating meta-cluster for traceability of lineages.

TrackSOM supports several modes of operation, related to the quantity and handling of meta-clusters. Under *Autonomous Adaptive* operation, the number of meta-clusters created at each time-point is automatically inferred. Alternatively, the user can specify the number of meta-clusters to be created as either a single number for all time-points/disease-stages (*Prescribed Invariant* operation) or a number for each time-point/disease-stage individually (*Prescribed Variant*). For each of these modes of operation, the merging of meta-clusters across time-points can be enabled or disabled. Disabling merging can inflate the number of meta-clusters above user-specified values, but better reflects the diverging nature of cellular differentiation.

### 2.2 TrackSOM tracks splitting, merging and transient clusters

We verified TrackSOM’s capacity to uncover cell population emergence, disappearance, splitting and merging in time-series data by applying it to a synthetic dataset into which these phenomena had been engineered. We manually gated the dataset to provide ‘ground truth’ labels of which data-points belong to the same population (Figure 2). TrackSOM performance was evaluated against these labels, using the Adjusted Rand Index (ARI) and tracking accuracy metrics for clustering and cluster tracking, respectively (Methods Section 5.5). To comprehensively assess TrackSOM’s performance we explored a broad range of parameter values. For each of TrackSOM’s three modes of operation, we generated 100 parameter value sets (Supplementary Material Table S2), varying the number of meta-clusters produced each day and the SOM grid size using Latin Hypercube sampling (McKay *et al*., 1979; Read *et al*., 2020). Meta-cluster merging was permitted.

**Figure 2:**
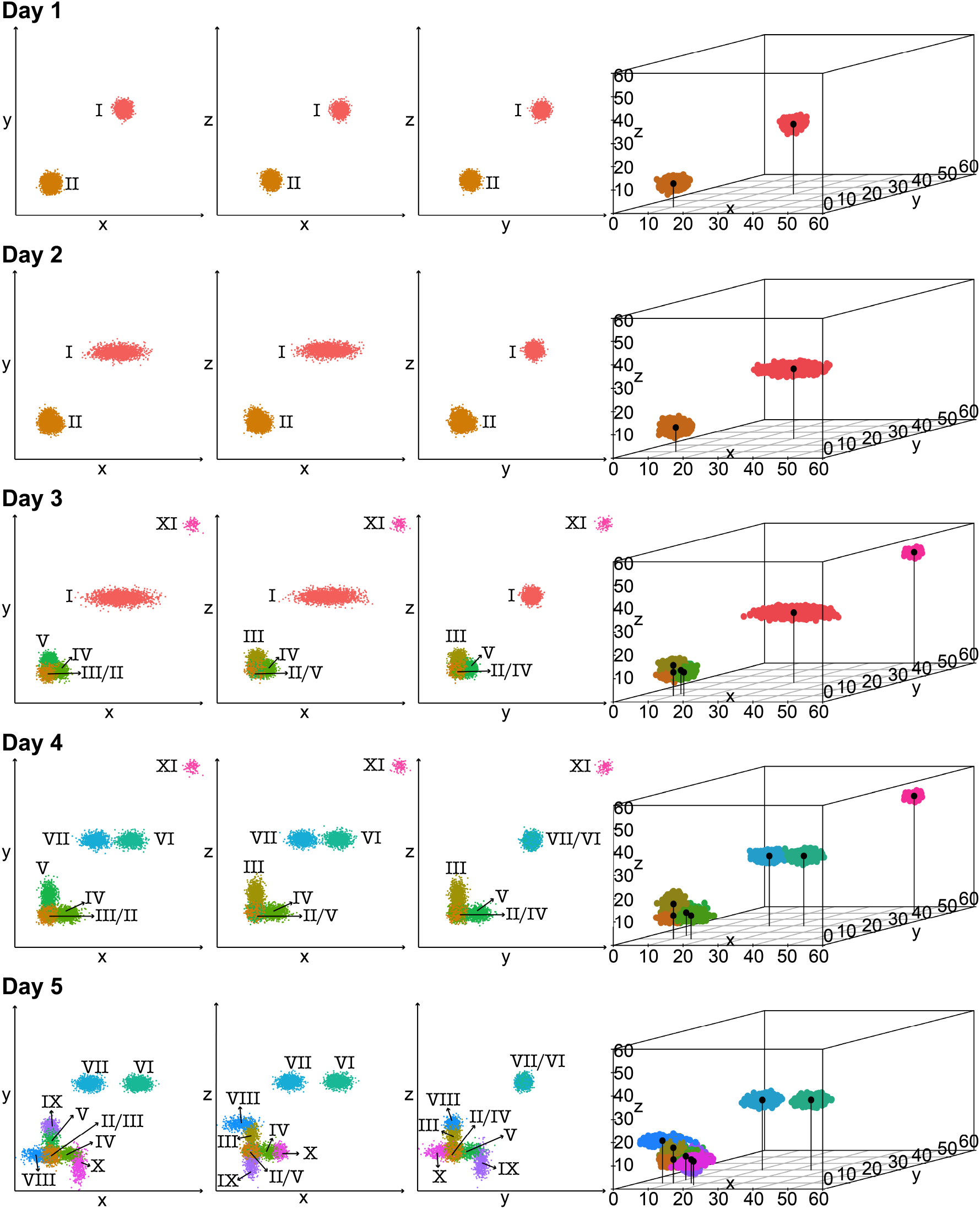
The synthetic dataset, shown as 2D (left) and 3D (right) scatter plots. Data-points are coloured by manual gating-assigned ground truth labels.

TrackSOM reproduced manually-gated populations and was robustly resistant to parametric perturbation. Solutions from *Prescribed Variant* operation consistently obtained mean ARI scores of at least 0.7 Figure 3A-B. TrackSOM’s other operational modes, *Autonomous Adaptive* and *Prescribed Invariant*, fared less well. They scored ~0.55 and 0.35-0.65 for ARI, respectively. *Autonomous Adaptive* operation was strikingly invariant to parameter value variation. Closer manual inspection of the clustered data (not shown) revealed it to consistently produce 3-4 meta-clusters per day and, consequently, a suboptimal capture of the sprouting conglomerate as a single meta-cluster. That *Prescribed Variant* operation obtained higher ARI scores than *Prescribed Invariant*likely reflects a better capture of the varying numbers of populations across time (Figure 2), to which *Prescribed Variant* can be adapted. Thus, overall, *Prescribed Variant* operation more accurately reflected both manually gated populations and their temporal evolutions.

**Figure 3:**
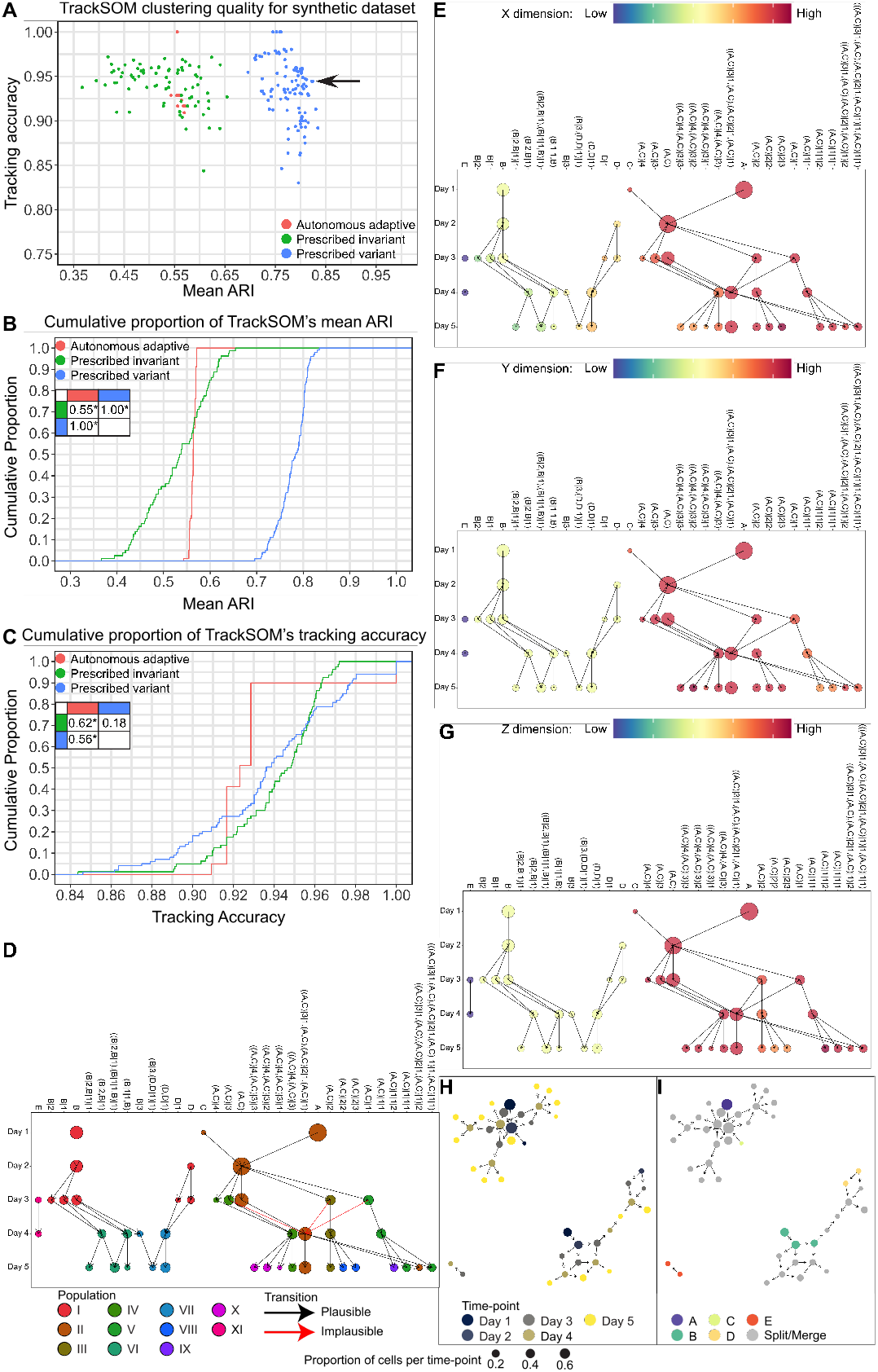
TrackSOM performance on the synthetic dataset. We generated up to 100 unique parameter value combinations for each of TrackSOM’s three modes of operation using Latin Hypercube sampling and then performed clustering on the synthetic dataset. Merging of meta-clusters was allowed. (A) A scatter plot of TrackSOM solution performances under the mean Adjusted Rand Index (ARI) over 5 time-points and tracking accuracy. We plotted the cumulative distributions of TrackSOM solution performances for (B) mean ARI and (C) tracking accuracy. Statistical comparisons are through the Kolmogorov-Smirnov (KS) test; * signifies p-value < 0.005. We selected a single high-performing TrackSOM solution to explore in greater detail (A, arrow). TrackSOM facilitates exploration of meta-cluster dynamics through several graphing formats. (D-G) Time-course heatmaps depict meta-clusters (dots), temporal evolutions (lines), relative meta-cluster sizes at each time-point (dot sizes), and either (D) the ground truth label it represents (see Fig. 3) or (E-G) the meta-cluster locations and movements through space (colours; each dimension shown independently). For (D), meta-clusters were assigned the ground truth label from which they captured the most data-points. Plausible and implausible transitions were determined based on the transition rules depicted in Figure S2. Network plots show meta-clusters coloured by (H) time-point and (I) cluster of origin (splits and merges are coloured grey). Arrows indicate meta-cluster evolutions over consecutive time-points.

TrackSOM excelled in correctly capturing temporal cluster dynamics, Figure 3A&C. Of the cluster evolutions uncovered, 83-100% reflected valid population transitions across the parameter perturbations explored. These were not simply self-referential transitions of the same populations across time-points; a quarter of the valid transitions were between distinct populations (e.g. meta-cluster (A,C), population II, splitting at day 2 to form meta-clusters representing populations III, IV and V, Figures 2D and S2).

To demonstrate data exploration through TrackSOM visualisations, we focused on one solution that performed well in both metrics (Figure 3A, arrow). This solution was generated under *Prescribed Variant*, generating 3, 3, 11, 9, and 17 meta-clusters per day from an 8×8 SOM grid. Between days 2 and 3, population II ‘sprouts’ into three different directions, each representing movements across X, Y or Z dimensions, Figure 2. This divergence is clearly captured by TrackSOM and shown in Figures 3E-G, where the branches stemming from meta-cluster (A,C) each move in only one dimension. Figures 3H-I allow meta-cluster lineages to be identified from their initial IDs, and show the slightly complex temporal transitions that have occurred, likely from a slight over-clustering of the data and having permitted meta-cluster merging.

### 2.3 Articulating the evolving immune response to West Nile virus infection in mouse bone marrow

We next evaluated TrackSOM against a real flow-cytometry dataset which captured the immune response in the bone marrows of West Nile virus (WNV)-infected mice over 8 days. The dataset was expertly pre-gated to generate ground truth labels from which ARI and tracking accuracy scores could be calculated. We executed each of TrackSOM’s three modes of operation both with and without meta-cluster merging, resulting in 6 operational modes in total. As with the synthetic dataset, we executed up to 100 unique parameter value combinations for each operation (Methods).

All of TrackSOM’s modes of operation were capable of generating high-quality clustering solutions. The majority of solutions scored over 0.7 for ARI, Figure 4A-B. *Autonomous Adaptive* operation was the least variable under parametric perturbation, but its ARI scores were sub-optimal owing to its persistent under-clustering of the data: it typically generated 8 meta-clusters despite manual gating defining 16 distinct cellular phenotypes (Figure S2). *Prescribed Variant* and *Prescribed Invariant* operations together generated the best ARI scores but were more sensitive to parametric perturbation with meta-cluster merging disabled. This is because under no-merging operation the number of meta-clusters specified is a strict lower bound and inappropriate parameter values can lead to an explosion in meta-cluster numbers, leading to reduced ARI scores. Conversely, with merging enabled, this parameter is not a strict lower bound and TrackSOM has greater capacity to correct for sub-optimal parameter values.

**Figure 4:**
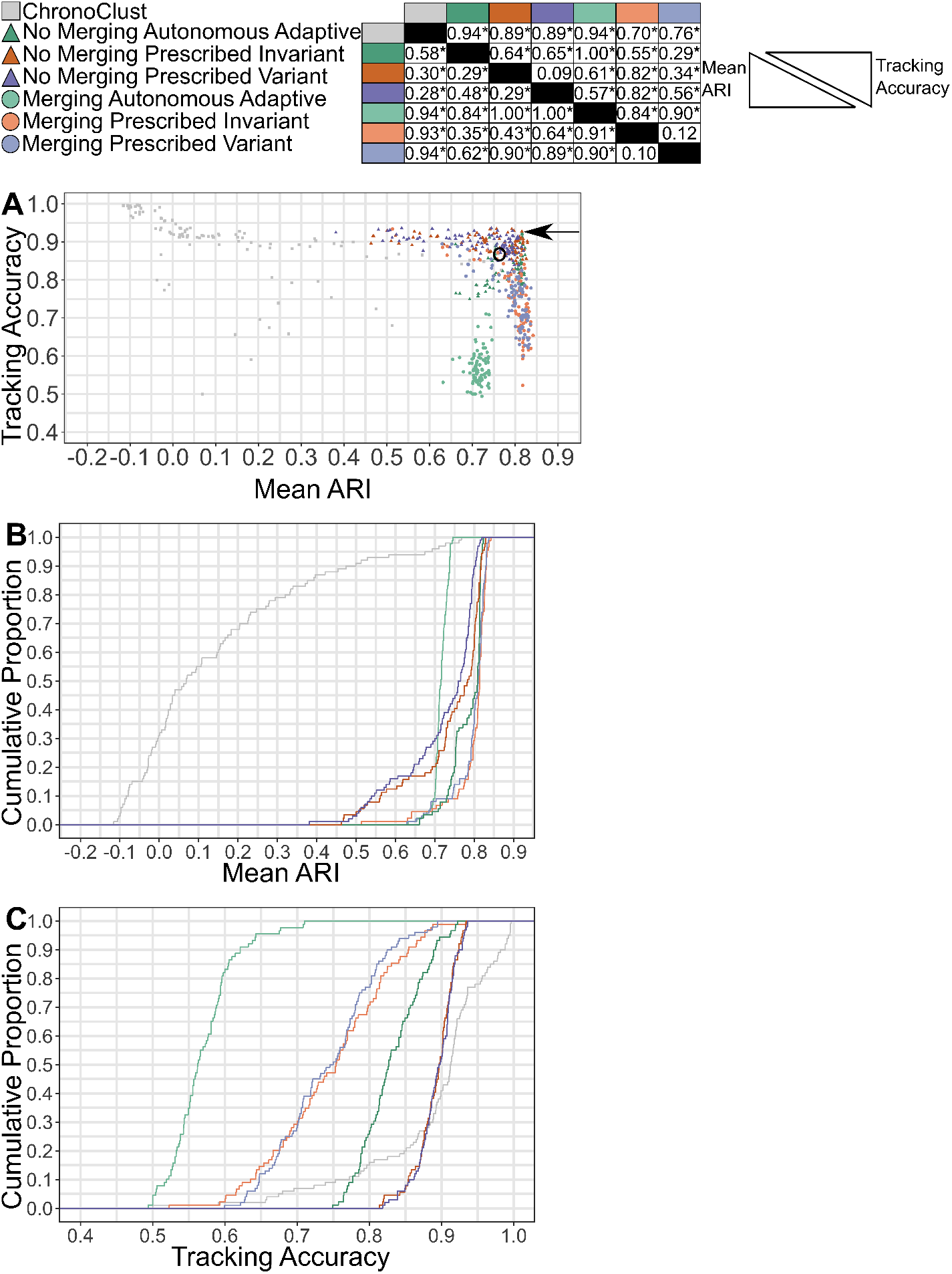
Evaluation of TrackSOM and ChronoClust clustering solutions when applied to the WNV bone marrow dataset. (A) Scatter plot of solutions in terms of mean clustering quality (ARI) over the 8 time-point dataset and the validity of the cluster temporal transitions generated (tracking accuracy). Solutions explored further in the text are denoted by the circle (ChronoClust) and the arrow (TrackSOM). (B-C) Cumulative distribution of TrackSOM's and ChronoClust's (B) Mean ARI and (C) tracking accuracy. Statistical comparisons for (A-C) are done through the Kolmogorov-Smirnov test.

TrackSOM performance in tracking evolutions of cell populations was more variable across its modes of operation, Figure 4A&C. *Prescribed Variant* and *Prescribed Invariant* were statistically indistinguishable and both out-performed *Autonomous Adaptive*, irrespective of whether merging was permitted. Interestingly, no-merging operation produced notably superior tracking accuracy scores than the same modes of operation when merging was permitted. This is because no-merging operation tended to generate more, homogeneous meta-clusters and self-referential transitions are valid (Figure S4). Conversely, merging operation has a greater capacity to amalgamate distinct cellular phenotypes and generate heterogeneous clusters, which can result in lower tracking accuracy scores.

Overall, the data reveal a possible trade-off between performance in clustering quality (mean ARI) and meta-cluster evolution tracking, as seen in the scatter plot of Figure 4A. The very best solutions by one metric are not necessarily optimal under the other. We were curious as to why TrackSOM exhibited such a wide range of tracking accuracy scores (0.6-0.9) at the leading edge of mean ARI scores (0.83). Mis-clustered data-points (leading to non-homogeneous meta-clusters) will suppress the ARI score to the same extent, irrespective of which meta-clusters they occur in. This is not the case for tracking accuracy, where all meta-clusters contribute equally to the score, irrespective of their size in terms of data-points. Mis-clusterings that are concentrated in a single, small meta-cluster at a given time-point could switch the assigned label of the meta-cluster and lead to invalid transitions, thus impacting tracking accuracy. If mis-clusterings are instead confined to large meta-clusters or broadly distributed across many meta-clusters, tracking accuracy is unlikely to be affected through ARI would be. Hence, the location of mis-clustered data-points could impact tracking accuracy whilst minimally affecting ARI. The explanation of why solutions could vary in mean ARI (0.38-0.83) whilst retaining high quality tracking accuracy (>0.9) lies in no-merging operation over-clustering the data and, as noted above, self-referential transitions are valid across time-points.

We selected one *No-merging Prescribed Invariant* solution that performed particularly well in both metrics for closer examination (arrow in Figure 5A). Figures 5A-B illustrate the solution as time-series heatmaps and network plots respectively. TrackSOM was able to track the progressive up-regulation of SCA-1 expression by several meta-clusters, and these mapped onto manually-gated cell populations transiting from un-activated to activated phenotypes. For instance, metaclusters E, E—1 and E—3 capture monocytes becoming activated at day 5, 3 and 5 respectively. Meta-cluster K represents stem and progenitor cells becoming activated at day 5, and metacluster G does likewise for T and NK cell populations. Not all the meta-clusters capturing a given population show up-regulation of SCA-1 marker: clusters P and K—1 also capture stem and progenitor cells yet show no such transition, demonstrating a capacity to uncover heterogeneity within phenotypes in response to infection. Further, being the bone marrow, activated forms of all the cell populations exist in small numbers even at pre-infection, and this is reflected in the cluster-population assignments, showing the sensitivity of TrackSOM to uncovering minor phenotypes in the data. For instance, meta-cluster N captures pre-infection activated B cells, and meta-cluster D does likewise for plasmacytoid dendritic cells (PDCs). The majority of cluster evolutions are biologically plausible (see Figure S2), although there are some exceptions, e.g. in meta-cluster M, B cells transit into T cells and NK cells. Such implausible transitions occur only in meta-clusters that capture a minority of data points (indicated by small symbols) and these likely reside on the periphery between several high-density regions of cell populations in marker-space.

**Figure 5:**
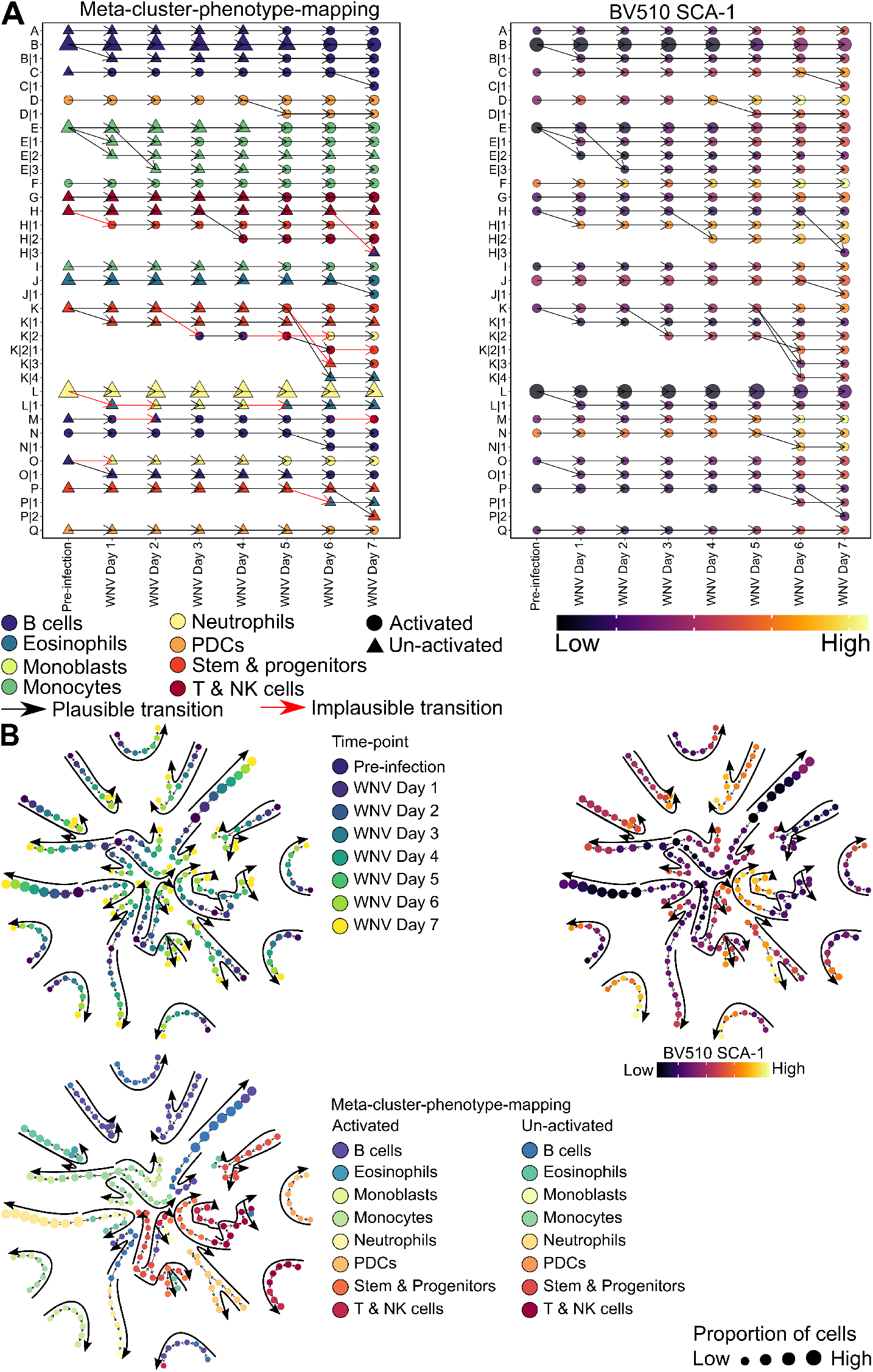
TrackSOM clustering of WNV bone marrow dataset using *No Merging Prescribed Invariant* operation. (A) Time-series heatmaps illustrating meta-clusters’ (symbols) temporal changes (lines). Meta-clusters on the left plot are coloured by the majority cell population they each represent. Biologically plausibility of transitions are indicated based on known cellular differentiation pathways (see Figure S2). The right plot captures average SCA-1 marker expression for each meta-cluster. (B) Network plots with meta-clusters coloured by time-point, cell population represented, and average SCA-1 marker expression.

Together, these data demonstrate that TrackSOM can accurately identify, without external guidance, both majority and minority population phenotypes and track their evolution through time in terms of changing marker expression levels. Further, it offers excellent performance in doing so without being overly sensitive to exact parameter value choices.

#### 2.3.1 Comparison with ChronoClust

We contrasted TrackSOM’s performance (above) with that of ChronoClust, the only other existing temporal cytometry clustering algorithm (Putri *et al*., 2019a), on the WNV bone marrow dataset. For a fair comparison with TrackSOM, we collected 100 unique parameter value combinations for ChonoClust (see Methods), and obtained their mean ARI and tracking accuracy scores.

All TrackSOM modes of operation vastly out-performed ChronoClust based on mean ARI score, Figure 4B. All TrackSOM solutions held ARI scores >0.38, whereas 30% of ChronoClust solutions scored <0, indicative of worse-than-random clustering. Curiously, these very poor ARI-scoring ChronoClust solutions also exhibited very high tracking accuracy scores (Figure 4A). Indeed, ChronoClust tended to generate higher tracking accuracy scores than TrackSOM, Figure 4C. Manual inspection of representative ChronoClust solutions revealed a pattern of severely over-clustering the data (Figure S4) into highly homogeneous clusters (poor ARI) that exhibited self-referential temporal transitions which are technically valid and thus promote high tracking accuracy scores. Examining Figure 4A more closely, the best ChronoClust solution (black circle on figure) held an ARI score of 0.76 and a tracking accuracy of 0.87. These scores are impressive, but still sub-optimal to what TrackSOM was readily able to produce in many solutions under parameteric perturbation. Indeed, the very wide distribution of ChronoClust solution scores suggests the algorithm to be quite sensitive to its parameter values, mirroring prior findings elsewhere Putri*et al*. (2021). Obtaining high-quality performance would require either considerable exploration of ChronoClust parameter space or prior knowledge of good parameter value choices. Overall, we consider TrackSOM the vastly superior algorithm, particularly under no-merging operation.

### 2.4 Advice on using TrackSOM

Key to the successful use of clustering algorithms on novel data is the selection of appropriate parameter values. We performed a global sensitivity analysis to relate TrackSOM’s two most influential parameters – SOM grid size and the number of meta-clusters generated – to its clustering and tracking performance. Accordingly, we calculated Partial Rank Correlation Coefficients (PRCC) of these parameters against TrackSOM performance on the WNV bone marrow dataset.

Examining the SOM grid size parameter first, Table 1, we find frequent positive correlations of grid size with mean ARI, yet negative correlations with tracking accuracy. For the *Prescribed* modes of operation, we find a non-linear diminishing increase in mean ARI as the SOM grid size increases (Supplementary Figures S8A, S9A, S10A, S11A). With SOM nodes being the fundamental unit of clustering, excessively small SOM grid sizes will capture distinct cellular phenotypes into the same nodes, ultimately reducing ARI scores. This effect was worse for no-merging operation, where small SOM grid sizes generate more variable and lower ARI scores (Figures S8A versus S9A), and S10A versus S11A, suggesting that merging operation has some capacity to correct for poor choices of SOM grid size. *Merging Autonomous Adaptive* operation exhibited an opposite pattern, with increasing SOM grid size reducing ARI scores, Figure S6A. We suspect this relates to an interaction between SOM grid size and how FlowSOM chooses the number of metaclusters to generate, see Supplementary Section Section S5.

**Table 1:** Partial Rank Correlation Coefficients (PRCC) of TrackSOM parameters against clustering (mean ARI) and tracking accuracy performance metrics for the indicated modes of operation. Data represents samplings of TrackSOM parameter space and corresponding clustering of the WNV bone marrow dataset. Only parameters with PRCC p-values < 0.005 are indicated (p-values not shown); ns, not significant.

| Operational mode | PRCC: Mean ARI | PRCC: Tracking |
| --- | --- | --- |
| **Parameter: SOM grid size** |  |  |
| Autonomous Adaptive, Merging | -0.72 | -0.50 |
| Autonomous Adaptive, No Merging | ns | ns |
| Prescribed Invariant, Merging | 0.62 | -0.93 |
| Prescribed Invariant, No Merging | 0.71 | -0.71 |
| Prescribed Variant, Merging | 0.78 | -0.95 |
| Prescribed Variant, No Merging | 0.82 | -0.66 |
| **Parameter: Maximum number of meta-clusters considered** |  |  |
| Autonomous Adaptive, Merging | 0.68 | ns |
| Autonomous Adaptive, No Merging | 0.38 | ns |
| **Parameter: Number of meta-clusters** |  |  |
| Prescribed Invariant, Merging | -0.55 | 0.88 |
| Prescribed Invariant, No Merging | -0.83 | ns |
| **Parameter: Number of meta-clusters per time-point** |  |  |
| Prescribed Variant (Pre-infection), Merging | -0.29 | ns |
| Prescribed Variant (Day 1), Merging | -0.31 | ns |
| Prescribed Variant (Day 2), Merging | -0.35 | ns |
| Prescribed Variant (Day 3), Merging | -0.39 | 0.52 |
| Prescribed Variant (Day 4), Merging | ns | 0.50 |
| Prescribed Variant (Day 6), Merging | ns | 0.32 |
| Prescribed Variant (Day 1), No Merging | -0.37 | ns |
| Prescribed Variant (Pre-infection), No Merging | -0.52 | ns |

There is a consistent pattern of increasing SOM grid sizes reducing the tracking accuracy. Tracking accuracy can diminish when the number of meta-clusters drops, because numerous meta-clusters tends towards self-referential transitions which are always valid. However, the presently observed pattern is independent of the number of meta-clusters created: PRCC controls for this parameter, and we find no discernible differences between merging and non-merging operations (the latter tends to raise meta-cluster numbers). This suggests that the pattern is driven by incorrect meta-cluster label re-assignments across time-points, which result in invalid population transitions. SOM nodes do not aim to capture equal numbers of data-points, instead they emphasise a comprehensive coverage of space, even between high-density regions. As such, increasing the SOM grid size will result in more nodes supported by fewer data points. Such nodes will be more sensitive to how the distribution of the data change between time-points, and for minority phenotypes (which are represented by relatively few nodes), these dynamics under large grid sizes could result in strong fluctuations of nodes entering and leaving the meta-cluster. We believe this could result in assignments of incorrect labels to meta-clusters and thus invalid transitions that reduce tracking accuracy scores.

As expected, increasing meta-cluster numbers tended to increase tracking accuracy for *Prescribed* modes of operation, though we found no effect for *Autonomous Adaptive* operation (Table 1, graphs in Figures S6-S11). This likely reflects increasingly homogenous meta-clusters, many of which will generate self-referential transitions as explored in the datasets above. For *Prescribed (In)variant* operational modes, increasing the number of meta-clusters reduced the ARI scores, likely due to the resultant over-clustering relative to manual-gated populations. Curiously, *Autonomous Adaptive* operation again exhibited a converse pattern, wherein the maximum meta-cluster number explored increased ARI scores; see Supplementary Section Section S5.

For users with un-analysed data we offer the following usage advice (Figure 6). If the number of cellular phenotypes to be discovered is completely unknown, *Autonomous Adaptive* operation can give a solid performance. *Prescribed Invariant* operation can be more effective, but is dependent on parameter values that such users are unlikely to have a basis to choose well; this is seen for both synthetic (Figure 3A-C) and bone marrow datasets (Figure 4A-C). *Autonomous Adaptive* operation can accommodate a large number of meta-clusters to explore generating (we attempted up to 40 with no deterioration in performance, Figure S6B & S7A), and we recommend users setting this to a large number as TrackSOM will adjust as needed. If the number of populations sought *is* known, then the *Prescribed (In)variant* operations are more appropriate. *Prescribed Variant* is more appropriate where the number of populations to be discovered varies over time, for instance via tissue infiltrates. If the number of populations is instead constant over time, then *Prescribed Invariant* is more appropriate and has fewer parameters to set.

**Figure 6:**
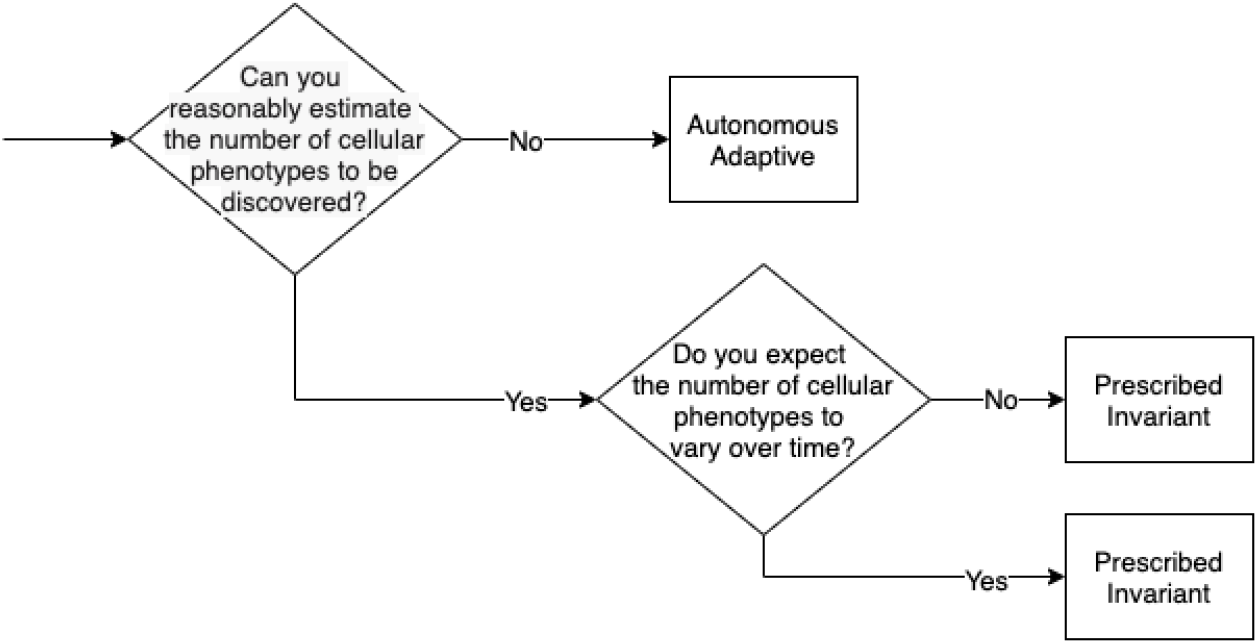
Practical guidelines for choosing TrackSOM’s mode of operation. As the separability and tracking of cellular phenotypes mostly depends on the number of meta-clusters for each time-point, the choice of operation mode is best guided by the user’s prior knowledge of the expected number of phenotypes in the data.

Preventing meta-cluster merging arguably better reflects the biology of cellular differentiation, and it conferred better performance in all operational modes, however this came at the possible expense of less meaningful clustering through an explosion of meta-cluster numbers if parameters are poorly chosen. However, the alternative - enabling merging - can instead generate convoluted cluster transitions. We suggest users try both modes and select that which better suits their circumstance. For selecting a SOM grid size, values between 10 and 15 consistently delivered a good balance between ARI and tracking accuracy scores (Figures S8-S11).

### 2.5 Mapping the CNS immune response onto WNV clinical disease severity

TrackSOM is not constrained to analysing time-course data, it can also relate immune system status to the spectrum of clinical disease severities that present at a single time-point. We demonstrated this in a novel, un-gated dataset representing the immune cells in the brains of WNV-(and mock-)infected mice at day 7 post-inoculation. A lethal dose (LD-100) was administered and mice were stratified into 6 groups based on the spectrum of clinical disease severities they exhibited at day 7, time of euthanisation. The corresponding cytometry datasets were organised into a sequence of increasing disease severities. We ran TrackSOM over this sequence to relate changes in cellular phenotypes to disease stage at this single time-point. In accordance with the above TrackSOM usage advice, we employed *Prescribed Variant* operation with meta-cluster merging disabled (see Methods) and conducted some manual exploration of meta-cluster and grid size numbers before manually selecting a solution that we judged to best separate the cell phenotypes. We anticipated that the number of cellular populations would increase with disease severity. This was supported by preliminary unguided analysis through FiTSNE, where increased severity associated with a larger number of distinct clusters of cells (Figure 7).

**Figure 7:**
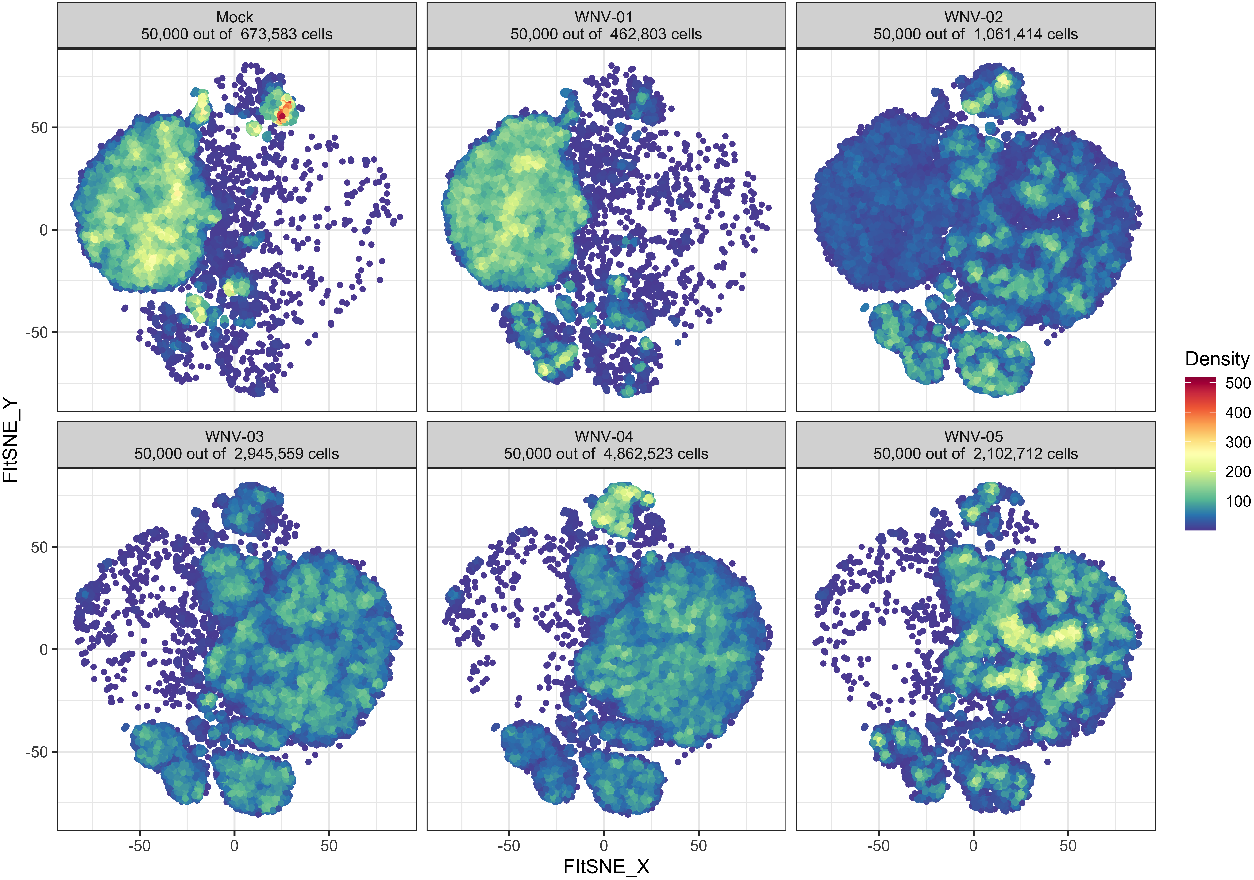
Progressing clinical disease severity seemingly associates with a shifting profile of immune cell populations in the brains of WNV-(and mock-)infected mice at 7 days post-inoculation. Shown are FiTSNE plots Linderman *et al*. (2019), stratified by disease severity (not time). Plots represent a subsampling of the total cells present at each disease severity, a pooling of these cells to calculate FiTSNE coordinates, and then plotting of cells by stage on each panel. Thus, populations are similarly spatially arranged across the panels. Cells are coloured by local density through kernel density estimation. Notably, the strongly represented cluster on the left side of the Mock dataset has not disappeared with increased disease severity, rather, those cells are diluted by a considerable increase in total cells present in the brains at later disease stages.

TrackSOM successfully distinguished numerous meta-clusters of cells, that we could resolve to immune cell phenotypes, and related their phenotypic evolution to increasing disease severity, Figure 8. TrackSOM tracked how the absolute (Supplementary Figure S12)and relative counts of cellular populations (Figures 9), and their marker expression levels (Figure 8 insets and grids) shifted with disease severity. As disease severity rose from mock-infection to the most severe WNV-05 we observed substantial rises in infiltrating macrophage, CD4 and CD8 T cell, NK cell and neutrophil populations. Further, we track a drop in the proportion of microglia from 80% to 15% of total cells, consistent with the concomitant substantial increases over this time in infiltrating macrophages. The proportion of infiltrating macrophages rose sharply from 10% to 70% of total cells by WNV-05. Several of these cell populations are not ordinarily found in the brain, yet were identified by TrackSOM in the mock-infection dataset, likely representing either the capture of post-flushing remnant intravascular cells in the samples and/or the recognition of central nervous system-associated macrophage populations in the homeostatic brain (Ivan *et al*., 2021).

**Figure 8:**
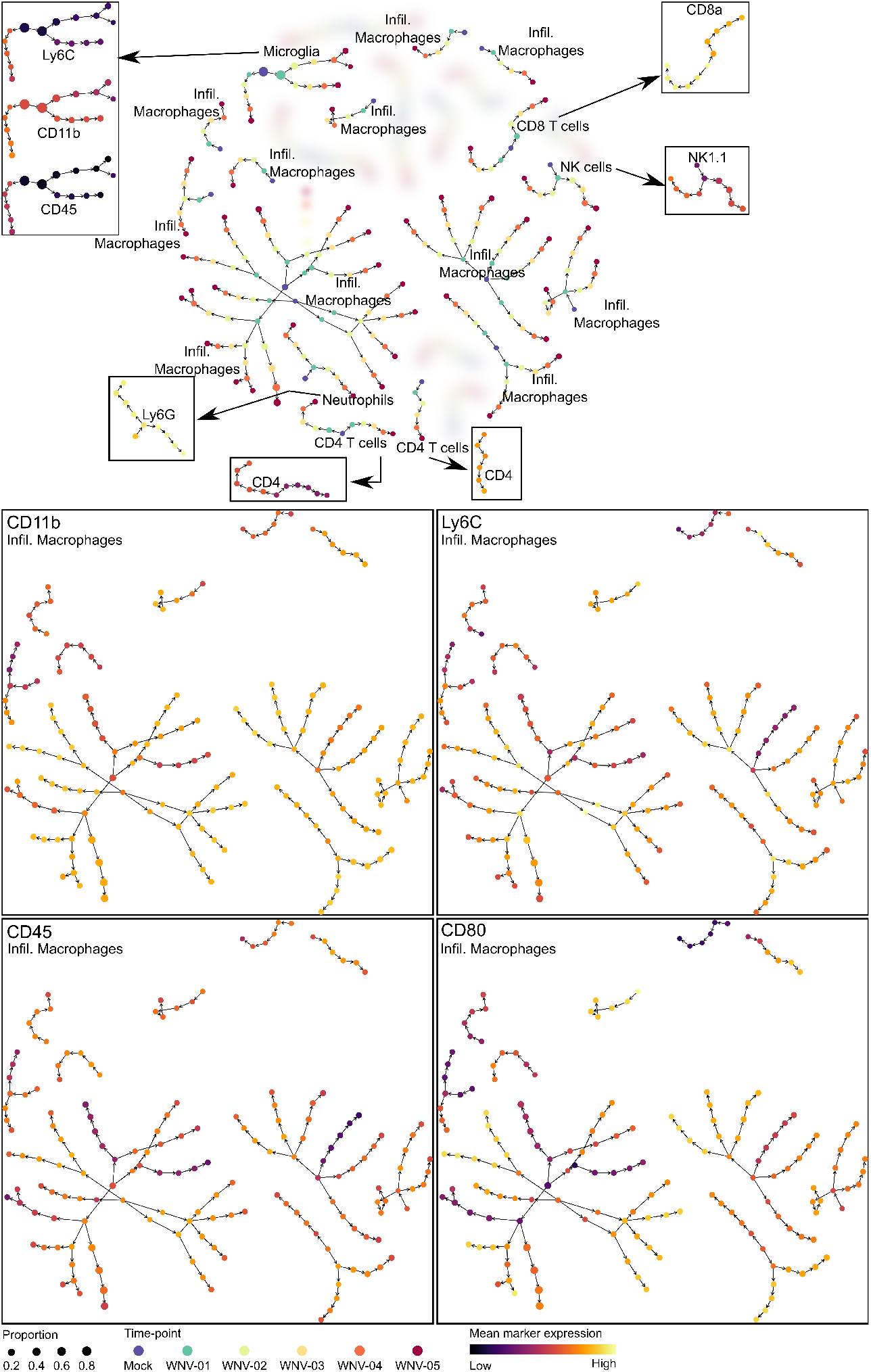
TrackSOM-enabled exploration of the immune response in the brains of WNV-infected mice over 6 graded clinical disease severities. All data represent day 7 post-inoculation. Shown is a network plot of TrackSOM meta-clusters and their evolution with worsening clinical disease severity at this time-point. Analysis is focused on meta-clusters given high-confidence manual phenotype labels (non-blurred). Meta-clusters in the central network plot are coloured by disease severity and arrows indicate the sequence of progression. Meta-clusters in the insets and grids are coloured by the indicated mean marker expression level. Node sizes relate to the proportion of cells captured in the meta-cluster at given disease severity. For insets, colourbars capture the full range of values for each marker independently.

**Figure 9:**
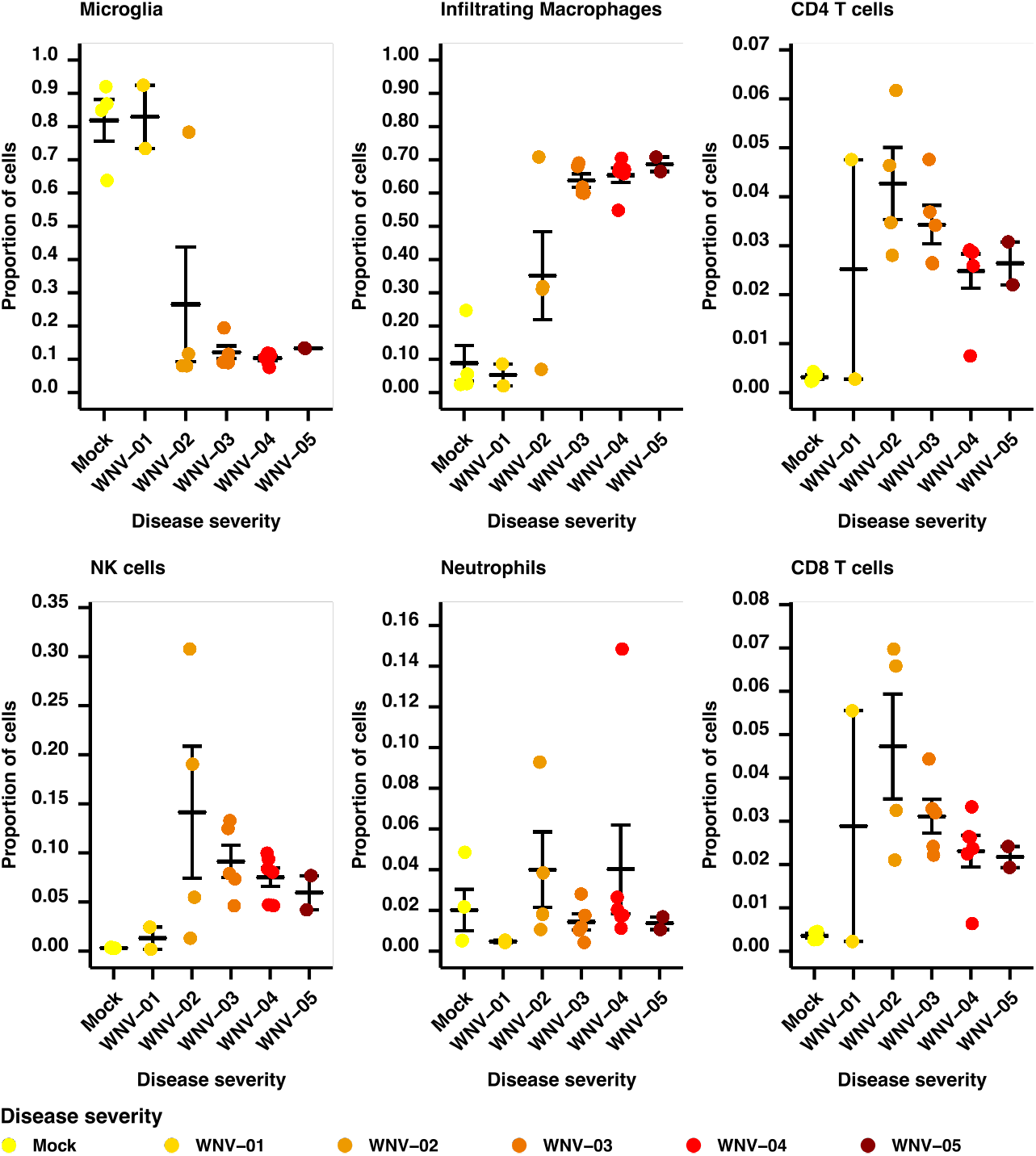
The profile of immune cell phenotypes varies with increasing disease severity in the brains of WNV-infected mice. Mapped are the proportion of the given cell phenotypes per mouse. Only cells (and TrackSOM meta-clusters) that we could label with high-confidence are included in the calculation of proportions. Data represent between n=2 and n=6 mice per group (Supplementary Table S5). Note that ranges for y-axes vary across panels.

Notably, TrackSOM identified the infiltration of phenotypic sub-populations with worsening disease severity, indicated by the branching meta-clusters in Figure 8. Macrophages were represented by 11 meta-clusters in mock-infection, curiously high given their low relative abundance (Figure 9A), and 4 of these underwent extensive branching into distinct sub-populations as disease progressed. Most of these macrophage sub-phenotypes up-regulated the CD80 activation marker with increasing disease severity, indicative of inflammation. NK cells were captured by one meta-cluster which upregulates the NK1.1 marker expression. Both CD4 and CD8 T cell populations were captured by meta-clusters that bifurcated at the point of mild (WNV-01) symptoms in accordance with CD4 and CD8 marker expression levels respectively, presumably capturing differentiating functions under immune challenge (Xiao *et al*., 2007). Microglia were captured by a single meta-cluster at mock infection that branched into several sub-lineages with heightening symptoms, varying in CD11b, Ly6C and CD45 expression.

Overall, TrackSOM was able to elucidate and map the changing profile of CNS immune cell populations onto a progression of clinical symptom severities emanating from WNV-infection.

## 3 Discussion

Immune responses are fallible. Identical inoculation doses can cause disease that resolves in some individuals but are lethal in others (King *et al*., 2007), and autoimmunity perhaps represents the quintessential immune system error (Li *et al*., 2018). In addition, many non-communicable diseases, of which prevalence rates are rising globally, have an immune system component (De Visser *et al*., 2006; Zeng *et al*., 2019). Many such diseases are characterised by periods of remission and relapse, often with localised exacerbation of inflammation, such as the skip legions that characterise Crohn’s disease (Roda *et al*., 2020) or the localised lesions in multiple sclerosis. The tone of the immune response, and its influence on health status, is a complex consequence of broad interactions between numerous dynamic cell populations (Li *et al*., 2018; Zeng *et al*., 2019).

Thus, the identification and characterisation of novel cell sub-populations has potential therapeutic benefit (Krzywinska *et al*., 2016). Mapping the immune response, at the level of cell population dynamics and interactions, over time and in relevant organs of the body, and relating these maps to disease outcomes, is essential for discovering possible intervention targets.

We developed TrackSOM with a view to enable mapping of the immune response against time and/or disease severity. TrackSOM fuses FlowSOM (Van Gassen *et al*., 2015), an effective, fast and widely-adopted automated gating algorithm, and ChronoClust (Putri *et al*., 2019a), a temporal clustering algorithm. TrackSOM excelled in unveiling the temporal evolution of clusters engineered into a synthetic dataset. When applied to pre-gated, time-series bone marrow cytometry data from WNV-infected mice, TrackSOM correctly tracked the activation of several immune cell populations and the up-regulation of the SCA-1 marker. On un-gated data, TrackSOM identified and tracked the infiltration and functional evolution of distinct (sub)populations of key immune cell phenotypes and mapped these against clinical disease stage. Further, TrackSOM readily revealed how the phenotypic composition, in terms of both absolute cell counts and relative abundance, varied with disease severity. TrackSOM is a powerful tool for studying coordinated immune population dynamics and how these pertain to disease status; it has already been used to identify cell populations and their shifting functional status over time in COVID-19 infection in humans (Koutsakos *et al*., 2021).

When evaluating clustering solutions, we observed a degree of trade-off across our two algorithm performance metrics. Solutions judged optimal for cluster quality tended to be poorer for cluster evolution tracking, and vice versa, particularly for Chronoclust. This is seemingly contradictory; how could poor quality clusters be well-tracked? The tracking accuracy metric is somewhat ignorant of cluster quality, and judges specifically only if cluster transitions are valid. Over-clustering, wherein single phenotypes are split across numerous meta-clusters, will tend to produce valid, self-referential transitions, thus scoring low on ARI but high on tracking accuracy. Under-clustering, wherein single meta-clusters amalgamate several phenotypes, will adopt and retain the majority population label, again valid for tracking accuracy but poor for ARI. There are advantages to multi-metric evaluation, as single metrics (and thus values) struggle to comprehensively describe all aspects of a complex phenomenon. Complementary perspectives through multiple metrics can offer a robust and comprehensive evaluation framework (Putri *et al*., 2021), and is the approach we took here. We identified and focused on solutions that exhibited minimal trade-offs between the two metrics, those solutions that generated an optimal balance of the conflicting criteria. With only two metrics, scatter plots such as Figures 3 & 4 were sufficient to identify such solutions. Where more metrics are employed, and the potential trade-offs more severe, a formalised framework through Pareto fronts can be employed (Putri *et al*., 2021).

TrackSOM’s *Autonomous Adaptive* operation carries considerable appeal, being largely insensitive to exact parameter value choices and thus simplifying the task of choosing appropriate values. However, we found it to consistently under-cluster the data, failing to fully distinguish all the subtly different cell populations discovered through expert manual gating. Its performance lagged behind the best that the *Prescribed (In)Variant* operations could produce when given optimal parameter values. *Autonomous Adaptive* operation draws upon FlowSOM’s procedure for self-determining an appropriate number of meta-clusters. The selection is made by identifying the point at which the reducing variance captured per meta-cluster sharply diminishes as the number of meta-clusters generated increases (Van Gassen *et al*., 2015) (a standard approach in clustering called the ‘elbow method’). FlowSOM’s method is simple and quick to execute, however it evidently does not converge on a clustering that an expert would manually produce. Identifying an alternative method that more closely aligns with expert opinion in cytometry specifically would be transformative for the discipline, enabling a heuristic search for appropriate parameter values that algorithm users must currently best-guess at and laboriously evaluate. This is a worthy avenue of future work, though prior efforts indicate that it is non-trivial (Wiwie *et al*., 2015).

A key investigative theme in immunology is to relate the kinetics of immune cell (sub)populations with disease outcomes, and to identify the key events at which outcomes bifurcate; these point to potential therapeutic targets. TrackSOM’s primary contribution is as an algorithm that automatically identifies and tracks the dynamics of cellular (sub)populations, mapping these over time and/or progressing disease status. A salient line of future investigation is the automated extraction of features in the rich data TrackSOM generates that constitute key distinguishers of clinical outcomes. For instance, TrackSOM could be run twice, over datasets of patients who recover versus those who do not. Within a supervised machine learning context, classifiers could be built over TrackSOM-originating features describing (sub)population kinetics to highlight those of particular relevance. Further, the classifiers would be of natural clinical benefit, potentially informing clinical decisions for novel patients by predicting their future clinical outcomes. Consider a contemporary possibility: predicting in advance which COVID-sufferers would require ventilators or admission into intensive care units. We are currently pursuing such technologies.

We focused here on cytometry because it is relatively cheap to run, accessible and widely adopted. Single-cell RNA sequencing (scRNAseq) is a related technology with falling costs and growing popularity. There is interest in identifying phenotype differentiation pathways through scRNAseq using ‘trajectory inference’ algorithms (Setty *et al*., 2016; Street *et al*., 2018; Trapnell *et al*., 2014; Saelens *et al*., 2019; Wolf *et al*., 2019; Van den Berge *et al*., 2020). This resembles TrackSOM’s usage context, and whilst trajectory inference algorithms have been applied over amalgamated time-series datasets (Byrnes *et al*., 2018; Scavuzzo *et al*., 2018), the algorithms themselves do not account for time, and thus do not relate changes to specific time-points. Pseudodynamics (Fischer *et al*., 2019) added an explicit accounting for time to trajectory inference methods. ScRNAseq differs from cytometry in that the data generated typically cover many thousands of features (genes) and hundreds of cells, whereas cytometry captures under 100 features and up to millions of cells. How well Pseudodynamics can accommodate cytometry data, and how well TrackSOM performs over scRNAseq data is worthy of future investigation.

## 4 Conclusions

The TrackSOM algorithm operates over cytometry data to automatically identify distinct cellular phenotypes and map their changing profiles onto progressing clinical disease stage and/or time. It provides critical support to analyse the immune response in terms of the dynamic and evolving, multi-faceted system that it is. TrackSOM was used to map out the immune response in the bone marrow over a week of time in WNV-infected in mice. TrackSOM outperformed ChronoClust, the only other existing temporal clustering and tracking cytometry algorithm, in terms of both quality of clusters generated and accuracy in tracking them. Making use of a comprehensive sensitivity analysis that provided putative users with operational advice, TrackSOM was applied to a non-pregated dataset to generate insight into how and when the phenotypic composition of immune cells in the brains of WNV-infected mice altered with worsening disease severity. In both brain and bone marrow, TrackSOM identified sub-populations within most phenotypes that exhibited differing trajectories through marker-space. TrackSOM is fast to execute and encompasses several modes of operation that can be chosen to best suit a given user’s experimental context. This includes a mode that requires very little parameter tuning to obtain high-performance results. The explicit tracking of cellular population dynamics provided by TrackSOM paves the way for integrated characterisation of the evolving immune response and mapping this to disease development and resolution.

## 5 Materials and Methods

### 5.1 TrackSOM algorithm

TrackSOM reads in time-course data in either FCS or CSV file formats. TrackSOM is written in R and uses FlowSOM’s R implementation as available from https://bioconductor.org/packages/release/bioc/html/FlowSOM.html. By default, prior to the start of building the SOM, TrackSOM independently normalises each feature across all time-points using the z-score normalisation provided by FlowSOM’s ReadInput function.

TrackSOM inherits FlowSOM’s parameters and their respective default values, though these default values can be changed to suit the user’s needs. Our present study focuses on varying the SOM grid size and the number of meta-clusters, parameters known to be most influential on FlowSOM clustering performance; all other parameters retain their default values. A full list of adjustable parameters is available in Supplementary Tables S1 and S2.

For *Autonomous Adaptive* operation the number of clusters trialled ranges from 3 to a user-specified maximum value. The value selected is that at which the rate of change in the variance captured in the meta-clusters suddenly decreases (this is adopted directly from the FlowSOM algorithm). For the *Prescribed (In)variant* modes of operation, the specified number(s) of meta-clusters is passed into FlowSOM’s consensus hierarchical clustering function for each time-point. If meta-cluster merging is disabled, user-specified meta-cluster numbers are a minimum value. FlowSOM-based meta-clustering of SOM nodes proceeds as usual, but TrackSOM will amend the result to split apart meta-clusters that would otherwise represent mergings from the preceding time-points.

For every cell, at every time-point, TrackSOM outputs which SOM the cell was captured by, which meta-cluster (if any) the cell was represented by, and that meta-cluster’s evolution through the dataset sequence. Together with the visualisations, this output allows users to associate metaclusters with cellular phenotypes. Network plots are arranged through force-directed layouts. The colouring of meta-cluster nodes by marker expression level on plots represents raw values mapped to the colour scales for each marker independently.

### 5.2 Synthetic dataset

The synthetic dataset is 3-dimensional (3D) and contains three distinct conglomerates of data-points that evolve over five time-points, Figure 2. We term them *conglomerates* as they encompass several Gaussian distributions that appear, disappear, move and/or change shape over time; these Gaussian distributions generate high-density sub-populations of data-points for TrackSOM to distinguish and track. The ‘sprouting’ conglomerate commences as a single high-density cloud of data-points that extend in three different directions over time. This mimics the transition of cells from common progenitors through different differentiation pathways. The ‘splitting’ conglomerate grows in two opposite directions, cleanly separating at day 4. The ‘transient’ conglomerate exists only at days 3 and 4. These two conglomerates mimic the infiltration of cell populations into the peripheral, challenged tissue, undergoing further differentiation and ultimately dissipating.

This dataset was adapted from that used in Putri *et al*. (2019a). It was generated by sampling data-points from non-isometric 3D Gaussian distributions that move, change shape, disappear and/or appear across five simulated time-points. Data-points are sampled afresh for each time-point - they do not persist between time-points. A total of 7000 data-points are generated for each time-point, with an additional 100 for days 3 and 4, representing a rare, transient cellular population. Manual-gating to allocate data-points into populations was performed by constructing non-overlapping polygons around the locations of Gaussian distributions. More information on the dataset’s specification, in terms of constituent Gaussian distributions, is available at https://github.com/ghar1821/TrackSOM.

### 5.3 WNV bone marrow dataset

This dataset has been previously reported in Putri *et al*. (2019a,b), and concerns quantifications of leukocytes in the bone marrows of mice over 8 days. Details on animal procedures are provided in Supplementary Section Section S3. The data comprise of 14 markers: FSC, SSC, Ly6C, CD45, CD48, Ly6G, CD117, SCA1, CD11b, CD11c, B220, CD115, CD16/32, CD3-CD19-NK1.1. Time-point 1 represents pre-infection (mock). Mice are infected with WNV on time-point 2 (WNV Day 1). There are 190,000 data-points (cells) per time-point. Author Ashhurst manually gated the dataset to assign each data-point (cell) a label representing its cellular phenotype, Supplementary Material Figure S1. TrackSOM was evaluated against these ground-truth labels. The following 16 cellular phenotypes were identified for each time-point, (1) Stem and progenitor cells, (2) B cells, (3) T and NK cells (indistinguishable given the markers in the flow panel), (4) Monoblasts, (5) Monocytes, (6) Eosinophils, (7) Neutrophils, (8) Plasmacytoid dendritic cells (PDC), with all 8 populations existing as activated or un-activated types. For assessing the tracking of temporal cluster dynamics, biologically valid transitions between phenotypes are known and shown in Figure S2, right.

The TrackSOM solution presented in Figure 5 was obtained using *no merging Prescribed Invariant* operation, creating at least 19 meta-clusters per day from a 7×7 SOM grid.

### 5.4 Algorithm parameter sweeps

We broadly explore TrackSOM and ChronoClust parameter space when clustering the WNV bone marrow dataset, and TrackSOM for the synthetic dataset. For TrackSOM, we generated 100 sets of parameter values using Latin Hypercube sampling for each dataset (McKay *et al*., 1979). For ChronoClust, we sub-sampled 100 results generated under Latin Hypercube from a prior publication using this dataset (Putri *et al*., 2019b). Briefly, Latin Hypercube sampling segregates each parameter’s range of values to be explored into discrete bins that are each sampled once and in a manner that minimises correlation between any two parameters. Hence, each parameter is partitioned into 100 bins, resulting in 100 unique parameter value sets. The strategy constitutes a broad yet efficient sampling of a parameter space (Read *et al*., 2020).

Latin Hypercube generates parameter values as floating-point numbers. However, because TrackSOM’s SOM grid size and number of meta-clusters parameters are encoded as integers, we round the parameter samples to the nearest integer before passing them to TrackSOM. This rounding can resolve to identical points in parameter space, and we discard duplicates from the analysis. Nonsensical parameter samplings, which specify more meta-clusters than the number of non-empty nodes available for any given time-point, were also discarded.

The parameters and ranges of values explored are given in Supplementary Material Table S2 and S3. The rationale for how parameter value ranges were chosen for each algorithm and dataset are provided in the Supplementary Material Sections Section S1 and Section S2.

### 5.5 Quantifying clustering and tracking performance

TrackSOM and ChronoClust performances were evaluated on datasets where ground-truth labels were available. The algorithms were executed over all available data-points in the dataset sequence, and evaluations were then performed on only those data-points (cells) for which ground-truth labels are available. This is standard practice in cytometry algorithm evaluation, as manual gating assigns only high-confidence labels to non-noise data-points (Weber and Robinson, 2016; Putri *et al*., 2021).

When applying TrackSOM and ChronoClust to the synthetic and WNV bone marrow datasets, for each time-point, each clustering solution was evaluated against the corresponding manually-gated ground-truth clustering using the Adjusted Rand Index (ARI) metric (Hubert and Arabie, 1985). ARI is a clustering similarity metric, representing the probability that two clustering solutions place any two randomly chosen data-points into the same cluster, and adjusting this for the rate of agreement reached under random assignment. ARI scores range from 1 to −1. The former represents a complete congruence in how two clustering solutions have clustered a dataset. A value of 0 represents the level of similarity expected if one of the clusterings represented random cluster assignment: some data-points of the same ground-truth label would still be clustered together. ARI values are obtained for each dataset in the sequence independently, and we then report the mean of these values.

To evaluate the quality of cluster evolution tracking, we adopted the tracking accuracy metric in Putri *et al*. (2019a). This metric computes the proportion of transitions which are valid (e.g., biologically plausible, such as un-activated B cells transitioning to activated B cells as characterised by up-regulation of the SCA-1 marker). Valid transitions for the synthetic and WNV bone marrow datasets are shown in Supplementary Figure S2.

The tracking accuracy metric requires each meta-cluster to be assigned a cell type (ground-truth) label. In our evaluation, each meta-cluster is assigned the majority label of the data-points it captured. In the event of a tie, alphabetical order of labels is used.

### 5.6 WNV CNS dataset

C57BL/6 female mice aged between 8 to 12 weeks were either inoculated or mock-inoculated with a lethal WNV dose (LD-100) and then culled for analysis 7 days later. Mice were stratified into 6 groups based on severity of symptoms at time of culling. As the WNV dose inoculated is universally lethal, the differential disease severity observed between groups at day 7 represent mice progressing through disease at different stages. Both the numbers of mice and the clinical scoring criteria are reported in Supplementary Table S5. Further details on animal procedures are provided in Supplementary Section Section S3. We applied TrackSOM in *Prescribed Variant no-merging* operation, specifying 25, 27, 30, 30, 30 and 30 meta-clusters be formed across the six groups from Mock to WNV-05. A 10×10 SOM grid size was used. We quantified the following 18 markers through flow cytometry: FSC, SSC, Ly6C, CD45, CD62L, CD4, CD86, CD11b, B220, Siglec-F, I-A/I-E, Ly6G, CD8-alpha, CD11c, CD115, CD80, CD3e, NK1.1.

### 5.7 Statistical methods

Latin Hypercube sampling and partial rank correlation coefficients (PRCC) were performed using the *Spartan* R package version 3.0.2 (Alden *et al*., 2013). PRCC values range from −1 to 1, indicating perfect monotonic negative and positive correlations respectively. The metric ascertains the effect of one parameter on TrackSOM performance whilst controlling for the effect of other parameters whose values will simultaneously be varying given the Latin hypercube design (Marino *et al*., 2008); this is a form of *global* sensitivity analysis (Read *et al*., 2012). We report only associations with *p* < 0.005, a value chosen to mitigate for multiple hypothesis testing of parameters against performance metrics across six modes of TrackSOM operation. ARI implementation was provided by the *mclust* R package version 5.4.7 (Scrucca *et al*., 2016). The FiTSNE plots and scatter plots for the WNV CNS data are provided through the *Spectre* R package version 0.4.0 (Ashhurst *et al*., 2021).

## 6 Declarations

### 6.1 Ethics approval and consent to participate

All procedures involving mice were reviewed and approved by the University of Sydney Animal Ethics Committee (AEC). The AEC fulfils all the requirements of the National Health and Medical Research Council (NHMRC) and the NSW State Government of Australia.

### 6.2 Consent for publication

Not applicable.

### 6.3 Availability of data and materials

TrackSOM is implemented in R, and is freely available to download under the GPL-3.0 open source license from https://github.com/ghar1821/TrackSOM. Code to reproduce all analyses and figures in the manuscript are also available at the aforementioned GitHub repository. All three datasets used here are freely available to download under the GPL-3.0 open source license from the Open Science Framework portal: (https://osf.io/8dvzu/).

### 6.6 Authors’ contributions

GHP, SD, IK, NJCK, TMA, MNR conceived the project. GHP, JC, DNE, FMW implemented the TrackSOM algorithm. TMA, performed the experimental work for the WNV datasets. GHP, TMA, MNR interpreted results. GHP, JC, MNR wrote the manuscript. SD, IK, NJCK, TMA, MNR supervised the project.

All authors read and approved the final manuscript.

## 6.7 Acknowledgements

We thank the Sydney Informatics Hub and Sydney Cytometry at the University of Sydney for providing access to their High-Performance Computing facility.

