## Supplementary Material for "TrackSOM: mapping immune response dynamics through sequential clustering of time- and disease-course single-cell cytometry data"

June 8, 2021

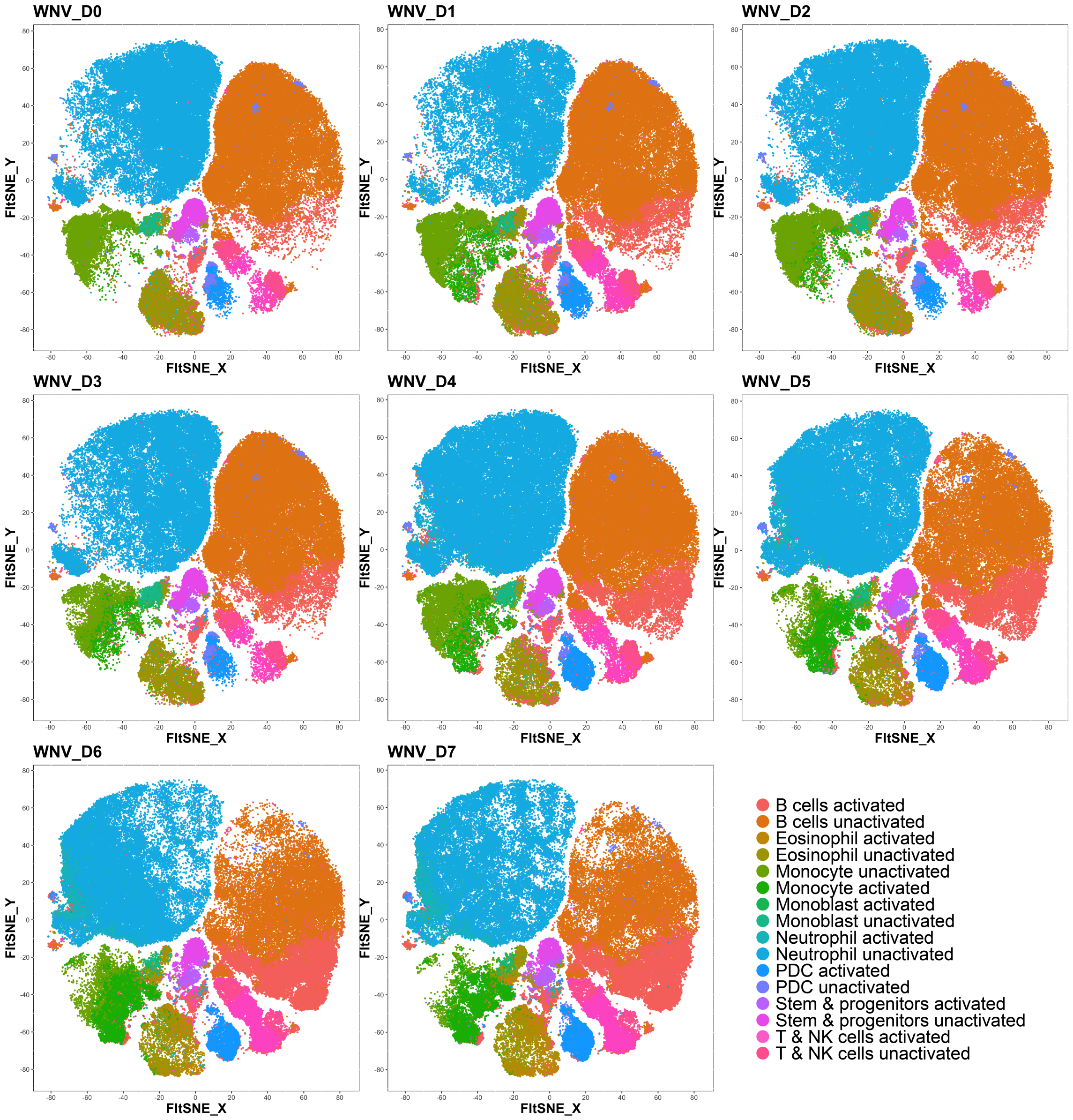


Figure S1: FIt-SNE [4] plots depicting the gated WNV bone marrow dataset. Each colour represents a manually-gated cell population. Data-points not gated into any population are excluded from the plot.

#### Section S1 TrackSOM parameter space sampling

For TrackSOM, Latin Hypercube sampling [5] was used to generate 100 putative samples of parameter space as floating point values. We varied the SOM grid size and the number of meta-clusters to be generated. As these parameters are encoded as integer values, some putative samples resolved to the same point in parameter space, and in this case duplicates were discarded. We retained: 100 unique parameter value combinations for *Prescribed Variant* operation, and 89 unique parameter value combinations each for *Autonomous Adaptive* and *Prescribed Invariant* operations. The remaining parameters (with exception to the random number seed which we set to 42) are set to the default values used in the FlowSOM implementation; Table S1.

The range of parameter values varied by Latin Hypercube is provided in Table S2. For all operations, we use a square grid size; rectangular grid sizes have not been found to confer any benefit [8].

| Parameter | Value |
|---|---|
| rlen | 10 |
| mst | 1 |
| alpha | 0.01-0.05 |
| radius | quantile(nhbrdist, 0.67) * [1, 0] |
| init | FALSE |
| initf | Initialize_KWSP |
| distf | 2 |
| silent | False |
| codes | Null |
| importance | Null |

Table S1: The values for TrackSOM parameters which were not varied under Latin hypercube sampling. These parameters are inherrited from the FlowSOM algorithm, and TrackSOM users can vary them if desired. FlowSOM's default random number seed is 32; in the present study we used 42 instead. See [8] and FlowSOM's vignette `https://bioconductor.org/packages/release/bioc/manuals/FlowSOM/man/FlowSOM.pdf` for more information on the parameters' functions.

| Mode | Parameter | Synthetic | WNV BM |
|---|---|---|---|
| Autonomous adaptive | Grid size | 5-15 | 7-20 |
| Autonomous adaptive | Maximum meta clusters | 5-20 | 16-40 |
| Prescribed invariant | Grid size | 5-15 | 7-20 |
| Prescribed invariant | Meta clusters per day | 5-20 | 16-40 |
| Prescribed variant | Grid size | 5-15 | 7-20 |
| Prescribed variant | Meta clusters per day | (3-4), (3-4), (6-12), (7-14), (9-18) | (16-40) per day |

Table S2: TrackSOM parameters varied under Latin Hypercube sampling. The ranges are applicable for both *merging* and *no merging* operations.

#### Section S1.1 TrackSOM parameter sweeps for the synthetic dataset

For autonomous adaptive, we chose to explore a range of 5-20 for the number of meta clusters per day. The minimum value is set based on the least number of populations we have per day (2 populations, day 1-2). As consensus clustering produces a minimum of 3 meta-clusters per day, to ensure *Autonomous Adaptive* operation is given a sensible range of meta-clusters to explore, we set the minimum number to 5. We set the maximum to 20, slightly over double the number of populations in the last time-point, to allow TrackSOM to better resolve populations which are small in size, such as the last appendages of the sprouting conglomerate in last time-point.

For *Prescribed Invariant* operation, we explored a range of 5-20 meta-clusters per day. The minimum of 5 meta-clusters is chosen to prevent severe under-clustering in latter time-points, in which we have at least 6 populations, while avoiding excessive over-clustering in earlier time-points where we only have 2 populations per time-point. The maximum of 20 is chosen to allow TrackSOM to better resolve the small populations in latter time-point.

For *Prescribed Variant* operation, a range of 3-4 meta clusters is chosen for first 2 days as TrackSOM produces minimum 3 meta clusters per day and that there are only 2 populations in those days. Setting the maximum to more than 4 will unnecessarily undermine the quality of meta-clusters for those days. On the subsequent days, we set the number of meta-clusters to be at least the number of populations expected, and at most double it.

For all operations, the SOM grid size is varied between 5-15. The value of 5 is chosen as the minimum as total SOM nodes must exceed the largest number of possible meta-clusters (20) in any time-point. The maximum value of 15 was selected so as to allow SOM nodes to transition between meta-clusters, thus reflecting the evolving nature of data in the input dataset sequence.

#### Section S1.2 TrackSOM parameter sweeps for the WNV bone marrow dataset

For the number of meta-clusters, we chose the same range for all operations: 16-40 meta-clusters per day. There are 16 manually-gated populations in the dataset, hence the lower value of 16 for this parameter. Our past experience (not shown) indicates that FlowSOM can better resolve populations when it is allowed to produce twice the number of meta-clusters as populations. Thus we set the maximum number for each time-point to be slightly more than double that of the 16 populations available.

For all operations, the SOM grid size is varied between 7 and 20. The minimum value of 7 is chosen to allow TrackSOM to produce 40 meta clusters for any given time-point. The maximum of 20 is chosen to allow TrackSOM to better resolve the small populations present in some time-points.

### Section S2 ChronoClust parameter space sampling

For ChronoClust, we varied all parameters except $\pi$. Parameter $\pi$ is defaulted to 14, which is the dimensionality of the WNV bone marrow dataset. Our prior experience (not shown) has revealed that ChronoClust tends to fail to produce any solutions (no data points are assigned to clusters) when $\pi$ is set to less than the dimensionality of the dataset.

| | Beta | Lambda | Epsilon | Mu | Delta | Upsilon | K | Omicron |
|---|---|---|---|---|---|---|---|---|
| Min | 0.1 | 0.125 | 0.01 | 0.0001 | 0.03 | 2 | 10 | 0.00001 |
| Max | 0.9 | 0.925 | 0.1 | 0.0009 | 0.1 | 10 | 20 | 0.0001 |
| Increment | 0.015 | 0.1 | 0.01 | 0.0001 | 0.01 | 1 | 1 | 0.00001 |

Table S3: ChronoClust parameters for WNV BM dataset. Pi is defaulted to 14.

Here, we randomly sub-sampled from the 400 parameter samples generated and used in [7], which spanned the range specified in Table S3, to extract 100 samples. Converse to the case with TrackSOM, Latin Hypercube does not produce any duplicate parameter space samplings for ChronoClust as it has float-encoded parameters.

### Section S3 Animal procedures

Female 9-10-week-old C57BL/6 mice were purchased from the Animal Resource Centre (ARC) (Western Australia, Australia). They were acclimatised over 1 week in individually ventilated cages under specific pathogen-free conditions with access to food and water ad libitum - in accordance with National Health and Medical Research Council's ethical guidelines. All experiments were completed with animal ethics approval by the University of Sydney Animal Ethics Committee.

Mice were anesthetised using isofluorane and infected intranasally with 1.2 x 105 plaque forming units (PFU) of West Nile virus (WNV), a dose that is lethal in 100% of mice without intervention, delivered in 10 $\mu$L of sterile PBS, as previously described in [3]. The original stock acquired from The John Curtin School of Medical Research (ACT, Australia) was propagated alternately in C57BL/6 suckling mouse brains and in vitro in Vero cells [2]. Mice were sacrificed no later than dpi 7m when they were anaesthetised and perfused transcardially with sterile phosphate-buffered saline (PBS). Brains and bone marrow were processed separately into single cell suspensions in PBS. Brains were processed in PBS containing DNase I (DN25, 0.05 mg/mL) and collagenase (5 mg/mL) (Sigma-Aldrich -MO, USA) using the gentleMACS dissociator (Miltenyi Biotec, Bergisch Gladbach, Germany). Subsequently, a 30%/80% Percoll gradient was used to isolate the leukocytes from brain homogenates. Bone marrow was collected from the bone marrow of the femur and tibia by flushing with PBS using a fine needle and syringe. Isolated cells were triturated to mechanically dissociate them into single cell suspensions in PBS without enzyme. After tissue

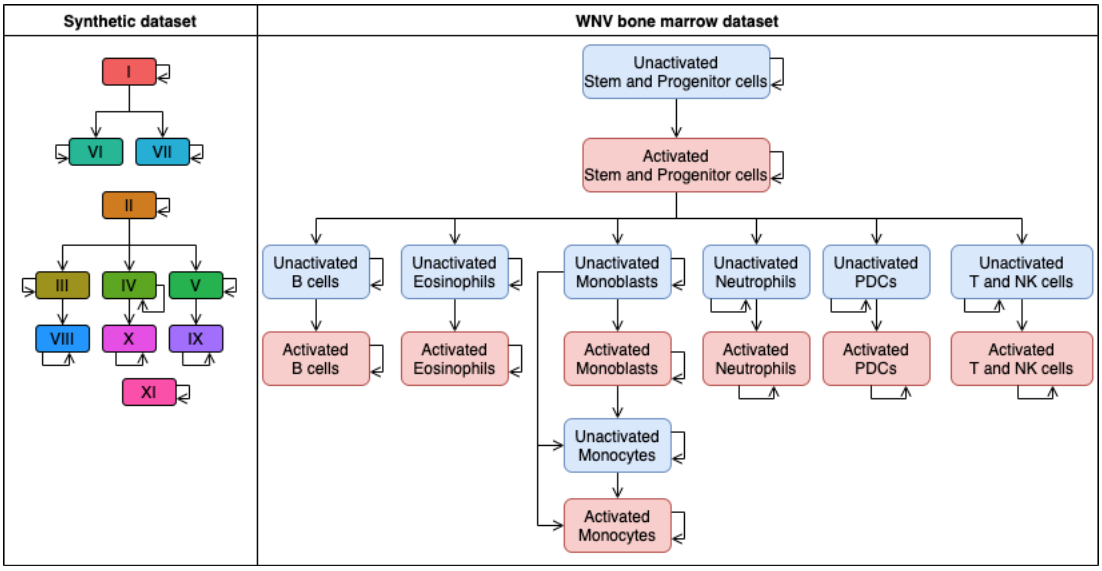


Figure S2: Valid temporal transitions between clusters, used in calculating the tracking accuracy scores. (Left) the synthetic dataset, and (right) the WNV bone marrow dataset.

processing, live cells were labelled with a specific panel of antibodies to define the relevant cell phenotypes [2, 1, 6].

### Section S4 WNV bone marrow dataset results

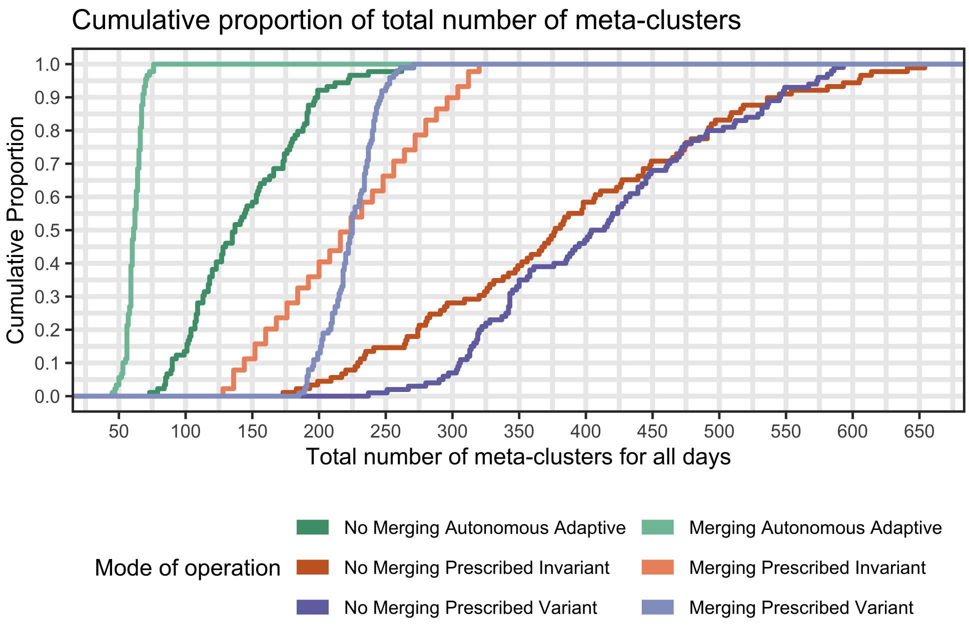


Figure S3: Cumulative proportion of the total number of meta-clusters generated by each solution under all TrackSOM's mode of operations when run on the WNV bone marrow dataset.

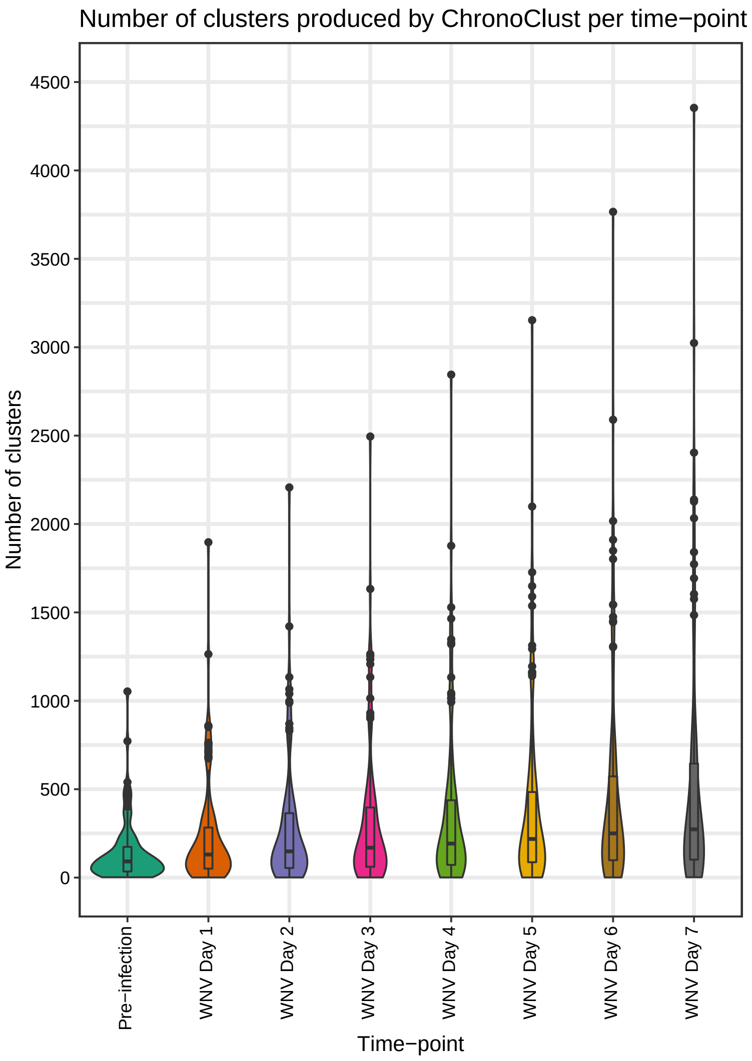


Figure S4: The number of clusters produced by ChronoClust for the WNV bone marrow dataset. There are 100 data-points for each time-point, representing the 100 parameter value samplings used in the analysis. Boxplots capture the median and interquartile range.

| | Number of clusters | | | |
|---|---|---|---|---|
| Time-point | Median | Mean | Minimum | Maximum |
| Pre-infection | 91.5 | 141.56 | 2 | 1,053 |
| WNV Day 1 | 130.5 | 233.65 | 1 | 1,897 |
| WNV Day 2 | 148.5 | 279.78 | 1 | 2,207 |
| WNV Day 3 | 169 | 315.42 | 1 | 2,495 |
| WNV Day 4 | 192.5 | 360.87 | 1 | 2,845 |
| WNV Day 5 | 218.5 | 407.25 | 1 | 3,153 |
| WNV Day 6 | 250 | 481.51 | 1 | 3,766 |
| WNV Day 7 | 273 | 537.77 | 2 | 4,354 |

Table S4: Median, mean, minimum, and maximum number of clusters generated by ChronoClust per time-point when run on the WNV bone marrow dataset. Statistics are representative of 100 ChronoClust parameter value samplings.

### Section S5 TrackSOM parameter sensitivity analysis, graphs and discussion

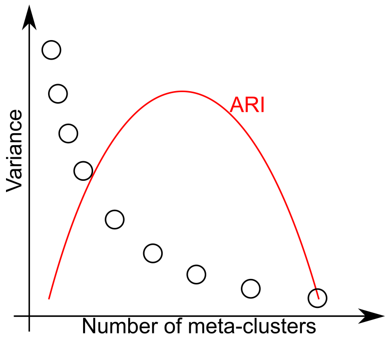


Figure S5: The relationships between the number of meta-clusters to be created, the variance in the data-points that a given meta-cluster captures (only one black dot shown per meta-cluster number) and the ARI scores (ref).

For *Merging Autonomous Adaptive* operation we found mean ARI scores to have 1) a negative correlation with SOM grid size and 2) a positive correlation with maximum number of meta-cluster considered. In this operational mode, FlowSOM infers the optimal number of meta-clusters based on an 'elbow criterion' calculation (see [8]). The underlying principle is that as the number of meta-clusters increases, the variance in the data they each capture will decrease at a diminishing rate (Figure S5). The 'elbow' is where the decreases start to flatten out, and the corresponding number of meta-clusters at this point is chosen. The calculation is based on two linear regressions, one on low and the other on high putative meta-cluster numbers. As the parameter representing the maximum number of meta-clusters to explore is increased, the slope of the second linear regression will become more shallow, and the elbow point will move to larger numbers of meta-clusters. The relationship of ARI to the number of meta-clusters is parabolic: too few meta-clusters represents an under-clustering wherein distinct phenotypes are pooled together; too many meta-clusters will split up what should be a single phenotype. With the values for the maximum meta-clusters to consider that we have explored, we believe FlowSOM is transitioning up the left side of the ARI parabola (Figure S5).

We are unsure exactly why SOM grid size correlates negatively with mean ARI score in *Merging Autonomous Adaptive* operation. We suspect that the additional nodes allow the data-points to be captured in a more granular manner, and that their collation into meta-clusters is resulting in meta-clusters with lower variance than for small SOM grids. This will impact the curve informing the 'elbow criterion' calculation, selecting a number of meta-clusters corresponding to a lower value on the ARI score parabola.

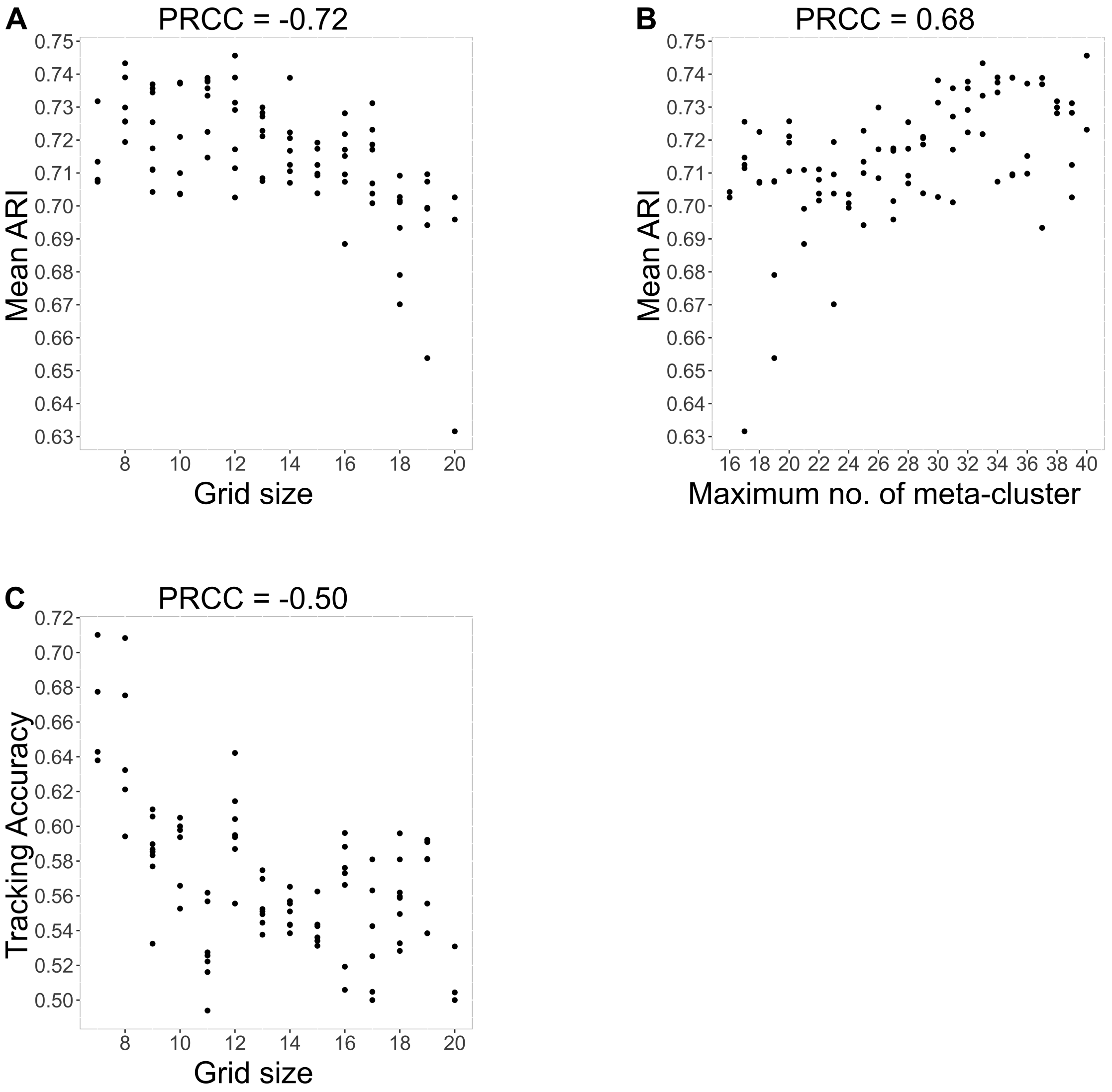


Figure S6: Scatter plots of TrackSOM parameter values versus clustering and tracking accuracy metrics, for *Merging Autonomous Adaptive* operation. Only statistically significant relationships are shown ($p < 0.005$). Partial Rank Correlation Coefficient (PRCC) scores indicate strength of associations.

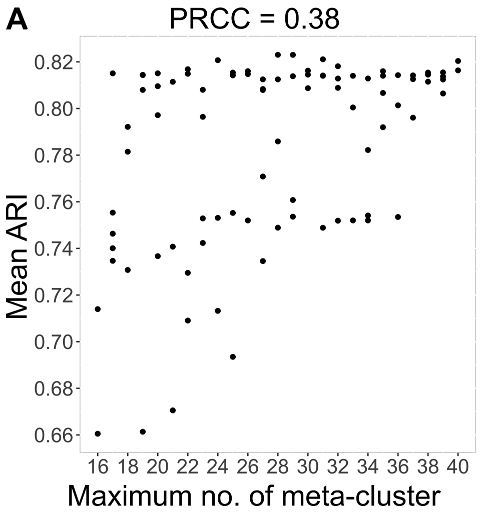


Figure S7: Scatter plots of TrackSOM parameter values versus clustering and tracking accuracy metrics, for *No Merging Autonomous Adaptive* operation. Only statistically significant relationships are shown (p < 0.005). Partial Rank Correlation Coefficient (PRCC) scores indicate strength of associations.

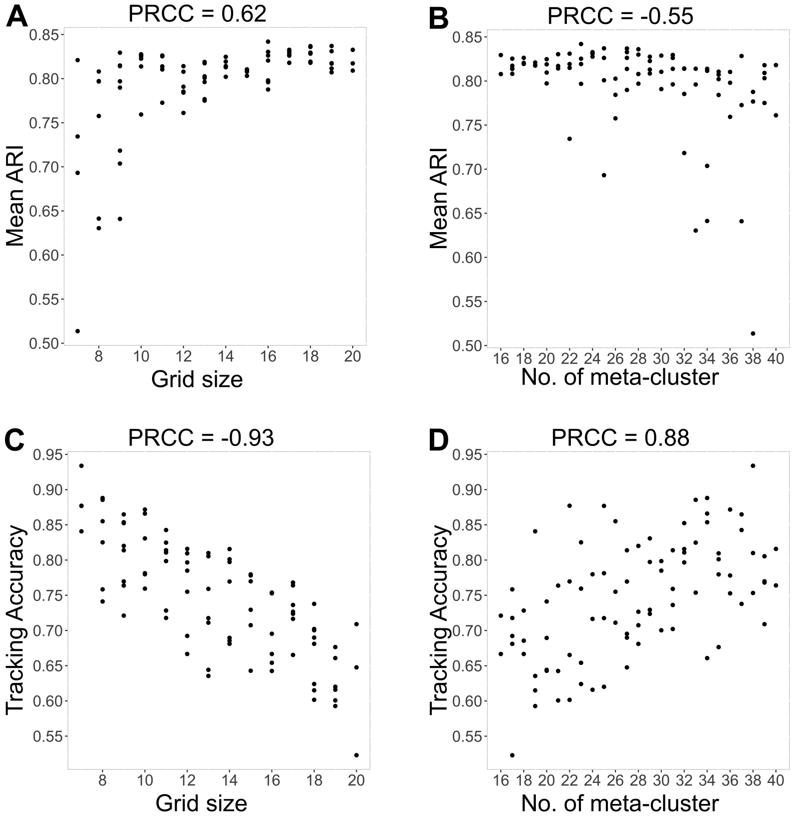


Figure S8: Scatter plots of TrackSOM parameter values versus clustering and tracking accuracy metrics, for *Merging Prescribed Invariant* operation. Only statistically significant relationships are shown (p < 0.005). Partial Rank Correlation Coefficient (PRCC) scores indicate strength of associations.

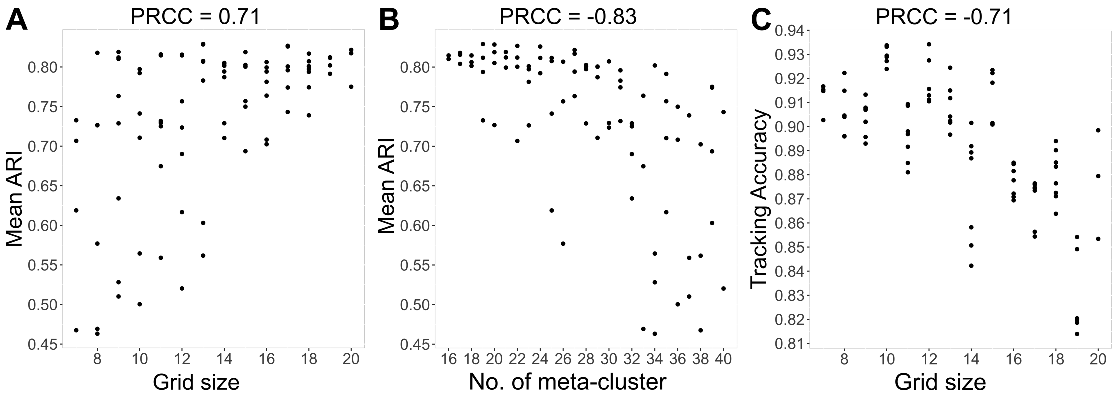


Figure S9: Scatter plots of TrackSOM parameter values versus clustering and tracking accuracy metrics, for *No Merging Prescribed Invariant* operation. Only statistically significant relationships are shown ($p < 0.005$). Partial Rank Correlation Coefficient (PRCC) scores indicate strength of associations.

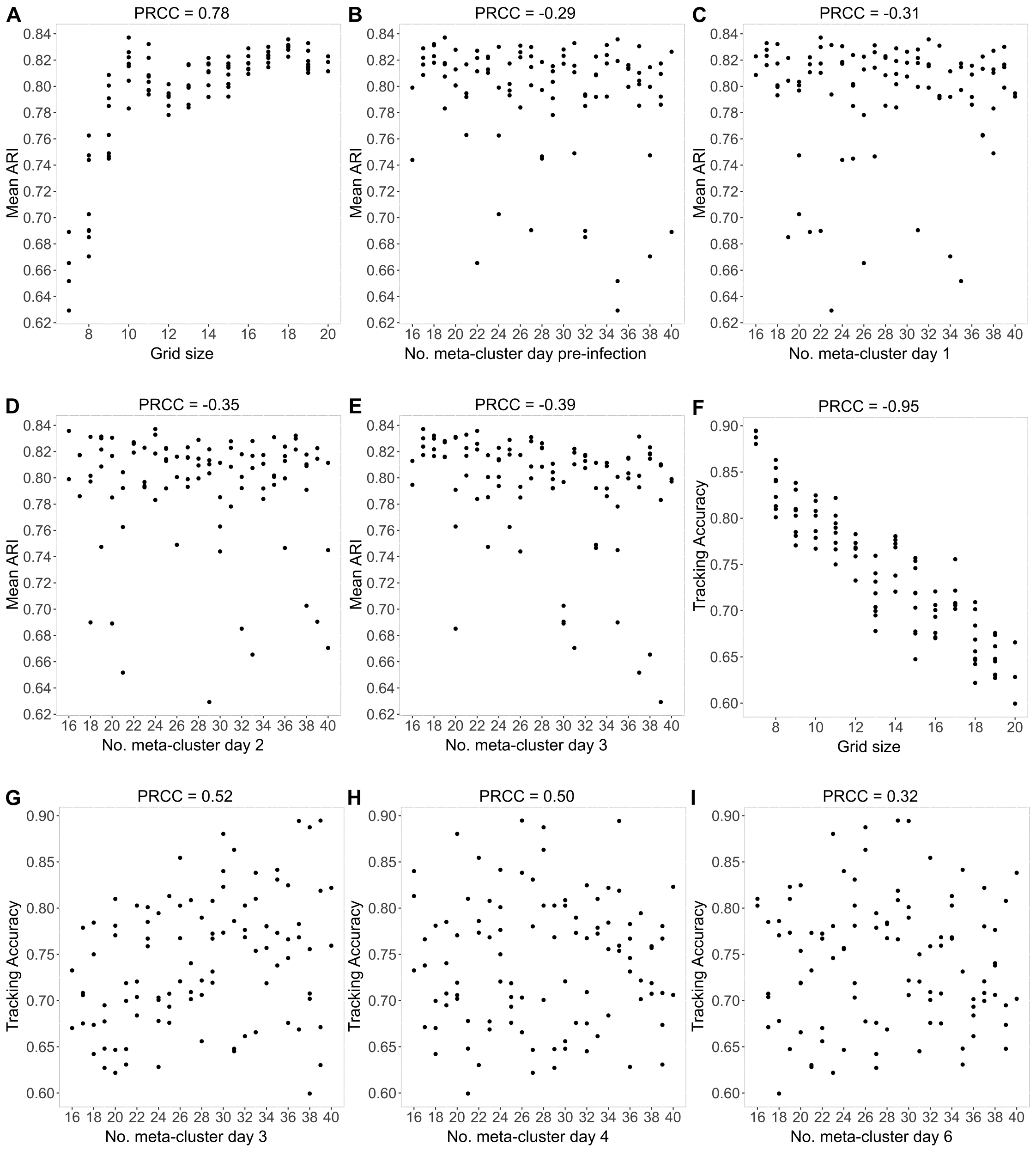


Figure S10: Scatter plots of TrackSOM parameter values versus clustering and tracking accuracy metrics, for *Merging Prescribed Variant* operation. Only statistically significant relationships are shown ($p < 0.005$) Partial Rank Correlation Coefficient (PRCC) scores indicate strength of associations.

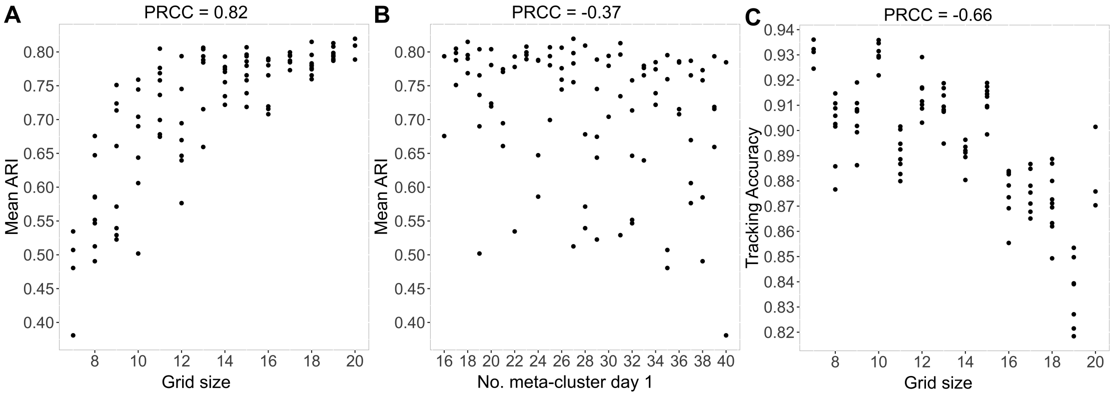


Figure S11: Scatter plots of TrackSOM parameter values versus clustering and tracking accuracy metrics, for *No Merging Prescribed Variant* operation. Only statistically significant relationships are shown ($p < 0.005$) Partial Rank Correlation Coefficient (PRCC) scores indicate strength of associations.

### Section S6 WNV CNS dataset

| Disease severity | Number of mice | Clinical criteria |
|---|---|---|
| Mock | 4 | Healthy mice (procedure but no inoculation). |
| WNV-01 | 2 | Minor slowing of movement. |
| WNV-02 | 4 | Slowed movement, but remaining responsive. |
| WNV-03 | 5 | Hunched, tendency to remain still, but able to move if prompted. |
| WNV-04 | 6 | Hunching, excessive face washing, rearing and falling, unwilling to move when prompted. |
| WNV-05 | 2 | laying on side and unresponsive, moribund. |

Table S5: Number of mice and clinical criteria for categorisation into each of the six disease severity groups in the WNV CNS dataset. Criteria were applied at day 7, at time of euthanasia for cytometry profiling.

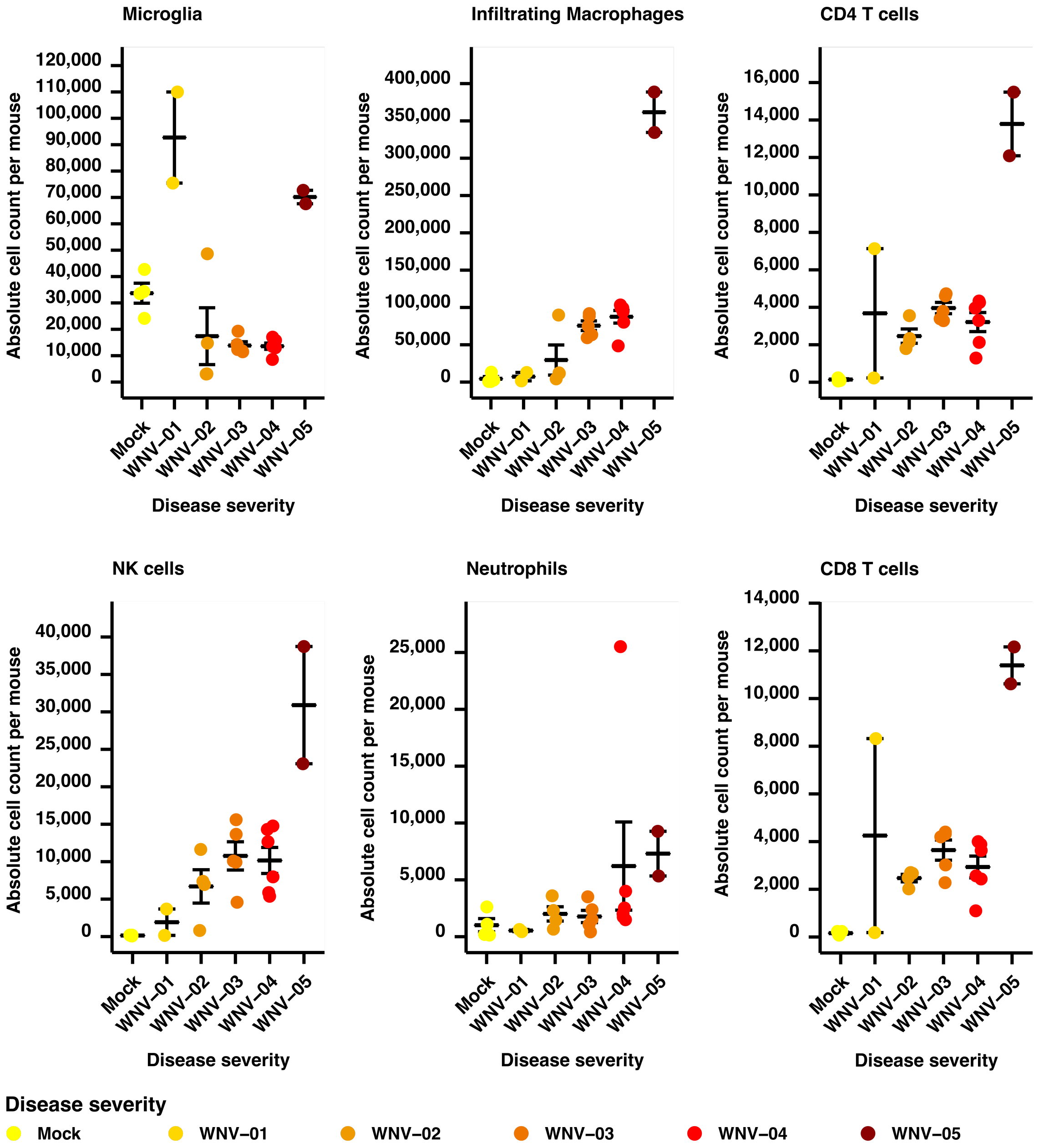


Figure S12: The profile of immune cell phenotypes varies with increasing disease severity in the brains of WNV-infected mice. Mapped are the absolute count of given phenotypes per mouse within each disease severity. Data represent between n=2 and n=6 mice per group (Supplementary Table S5). Note that ranges for y-axes vary across panels.
